# α-catenin phosphorylation is actomyosin-sensitive and required for epithelial barrier functions through Afadin

**DOI:** 10.1101/2025.08.21.671625

**Authors:** Jeanne M. Quinn, Phuong M. Le, Anthea Weng, Annette S. Flozak, S. Sai Folmsbee, Erik Arroyo-Colon, Mitsu Ikura, Noboru Ishiyama, Cara J. Gottardi

## Abstract

Zonula adherens junctions (zAJ) are spatially proximal to tight junctions (TJ), in a superstructure known as the apical junctional complex (AJC). A key component of the AJC is a circumferential ring of filamentous (F)-actin, but how actomyosin contractility drives AJC structure and epithelial barrier function is incompletely understood. Here, we show that a central mechanosensitive component of zAJ, α-catenin (α-cat), undergoes force-dependent phosphorylation in an unstructured linker region. This modification in turn primes the α-cat mechanosensitive Middle-region for effector-binding. We credential Afadin, a multi-domain TJ/AJ scaffold protein, as mechano-chemical binding partner of α-cat, identifying residues in α-cat required for this interaction. α-cat phosphorylation and Afadin-binding are required for their co-enrichment at zAJ and epithelial barrier function. A mouse model that prevents α-cat phosphorylation is particularly detrimental to post-natal brain development. These data support a stepwise model where α-cat integrates mechanical and chemical signals to progressively promote zAJ enrichment, effector recruitment and epithelial barrier function.

## INTRODUCTION

Zonula adherens junctions (zAJ) are a cell-cell adhesive structure long known to be spatially proximal to tight junctions (TJ), which control the passage of ions and small molecules between apical lumen and basolateral tissue compartments. Juxtaposed, these junction structures form a superstructure commonly known as the apical junctional complex (AJC)^1^. At both electron microscopic and super-resolution levels, these junctions appear compacted and invariantly aligned, with the TJ apical to zAJ^2,3^. A key component of the AJC is a circumferential ring of filamentous (F)-actin and myosin, where actomyosin contractility critically contributes to AJC structure, epithelial monolayer coordination and barrier function^4,5^. How actomyosin networks work with key protein constituents of TJ and zAJ to drive this universal organization of epithelial apical junctions remains incompletely understood.

Decades of research have identified major constituents of TJs and zAJs. Tight junctions rely on actin-binding scaffold ZO proteins to link multi-pass transmembrane spanning claudins, occludin/tricellulin and single-pass adhesive JAM proteins to the underlying actin cytoskeleton^1,6–8^. zAJ rely on two distinct actin-binding scaffold/transmembrane adhesion-receptor complexes: Afadin-Nectin and α-catenin/β-catenin-cadherin complexes, where the latter is critical for basic cell-cell cohesion and the former (via Afadin) for reinforcing junctions under dynamic cell-cell rearrangements and morphogenetic movements^9–14^. Recent super-resolution imaging of simple epithelial cell cultures may provide a clue regarding these functional differences. For example, Afadin-nectins localize to the apical-most portion of zAJ, apical to catenin/cadherin proteins and basal to TJ proteins in an intestinal epithelial cell culture model^3^. Evidence that F-actin abundance appears greatest at the level of Afadin-nectin complexes, where Afadin KO cells shows reduced junction alignment of F-actin ^15–17^ and greater apical membrane area due to lower apical tension^3^, suggests that Afadin plays a key role in supporting an apical actomyosin contraction apparatus that supports normal epithelial monolayer organization^3,18,19^. Since these junctional complexes mature and separate over time^3^, a set of rules must be at play to drive this process.

One emerging principle of junction biology is that mechanosensitive scaffold proteins use actomyosin forces to unfold and recruit partners required for zAJ organization and function^20–24^. α-catenin (α-cat) is one of the best known mechanosensitive proteins at zAJ, where it links cadherin adhesion receptors to the underlying actin cytoskeleton through a series of bundled α-helical domains that flexibly pin α-cat’s N-terminus to cadherins via β-catenin and C-terminal region to F-actin^25,26^. When myosin-generated forces pull on actin filaments, tension adjusts α-cat’s 5-helical actin-binding domain (ABD) to favor high affinity binding to F-actin^27–31^. This enhanced actin-binding allows forces to override salt-bridges that normally pin α-cat’s middle (M1-M2-M3)-region into a closed conformation. As a result, junctional forces can sequentially unfold α-cat’s M-region, with M1 unfolding at lower, followed by M2-M3 at higher forces^32–39^. Since the M-region is a major effector-binding region of α-cat^40–44^, α-cat is reasoned to link actomyosin force thresholds to distinct unfolded states and partner recruitment. But mechanosensitive binding sites for all but one α-cat effector (i.e, vinculin)^32,39,45–51^, remain poorly defined, and will be critical to matching individual effector binding roles to junction organization and functional reinforcement.

Afadin is a multi-domain scaffold protein that relies on Rap1/2-GTPase binding^12,52–54^, PDZ-membrane protein recruitment^9,55,56^, dimerization^57^ and intrinsically disordered and actin-binding domains^58,59^ to support AJ structure and dynamic regulation. Within its large intrinsically disordered region, Afadin also contains a coiled-coil (CC)-region (amino acids 1386-1595) that can bind the M3 domain of α-cat^15,41,43^, where this region contributes to Afadin enrichment and tight alignment of cortical actin at zAJ^15–17^. Whether effects of Afadin’s coiled-coil region are through direct binding to α-cat remains unknown, due to incomplete mapping of the Afadin-binding site on α-cat. Also unknown is the degree to which an α-cat/Afadin interaction depends purely on mechanical unfolding of α-cat, as opposed to other chemical signaling events. Lastly, whether mechanochemical regulation of this interaction is critical for epithelial barrier function in developing tissues remains untested.

Here we identify residues in the α-cat M3 domain required for Afadin binding, zAJ recruitment and epithelial barrier function. We show that Afadin-binding to α-cat requires an unfolded M3 domain, where phosphorylation of a linker-region proximal to M3 primes α-cat for M-domain unfolding and Afadin recruitment. Evidence that α-cat phosphorylations are actomyosin sensitive, sufficient to drive enrichment of α-cat and Afadin at zAJs, and required for organism viability suggests that junction scaffold proteins must integrate mechanical and chemical signaling inputs for epithelial barrier integrity.

## RESULTS

### Phospho-α-cat enriches at apical junctions and is actomyosin-sensitive

α-cat is highly phosphorylated in an unstructured region that links its mechanosensitive middle- and actin-binding domains^60–66^. We previously showed that this α-cat phospho-linker (P-linker) region is modified sequentially through a dual-kinase relay mechanism, where CK2 phosphorylates α-cat at S641, followed by sequential CK1 phosphorylations at S652, S655 and T658^66^. While modification of all these sites contributes to cell-cell cohesive strength, the overall logic of this rather elaborate dual-kinase mechanism remains a puzzle. We recently discovered that α-cat phosphorylation at CK1 triplet sites becomes elevated during mitotic rounding in HeLa cell lysates^67^. However, phospho-specific antibodies to these sites (pS652, pS655 and pT658) labeled all junctions in Madin Darby Canine Kidney (MDCK) cells^67^ (Fig. 1A), with no apparent enhancement between mitotic cell-neighbor junctions (Fig. 1B). These data suggest that α-cat phosphorylation may be more related to a cellular feature or property common to mitotic HeLa cells and MDCK cell monolayers^67^. A clue to understanding this difference was revealed by confocal optical sectioning in the x-z direction, where α-cat phospho-detection appeared biased towards the apical-most region of cell-cell contacts compared to an antibody recognizing total α-cat (Fig. 1C). Since the apical region of cell contact comprises closely aligned tight junctions, adherens junctions and a zonular belt of F-actin and myosin (hereafter referred to as zonular adherens junctions, or zAJ)^4,45,68–70^, we speculated that α-cat phosphorylation might be promoted by actomyosin contractility, a property that would be shared between mitotic HeLa cells and epithelia with prominent intercellular junctions. Indeed, α-cat phosphorylation at CK1 sites is diminished in cells treated with the myosin inhibitor, Blebbistatin (Fig. 2A). Conversely, α-cat phosphorylation at these same sites is enhanced by the actomyosin contractility promoter and myosin phosphatase inhibitor, Calyculin (Fig. 2B-C). The impact of these inhibitors is also seen acutely if CK1 phosphorylation is first inhibited and actomyosin regulators are present during the wash-out step (Fig. 2D-E). Under these conditions, the actin polymerization inhibitor, Latrunculin A, also attenuates α-cat phosphorylation. Together, these data suggest actomyosin contractility favors α-cat P-linker phosphorylation at its CK1 sites, and are consistent with previous evidence these sites are less accessible to *in vitro* modification of full-length α-cat relative to a fragment lacking N-terminal sequences^66^ (Fig. 2F).

**Figure 1:**
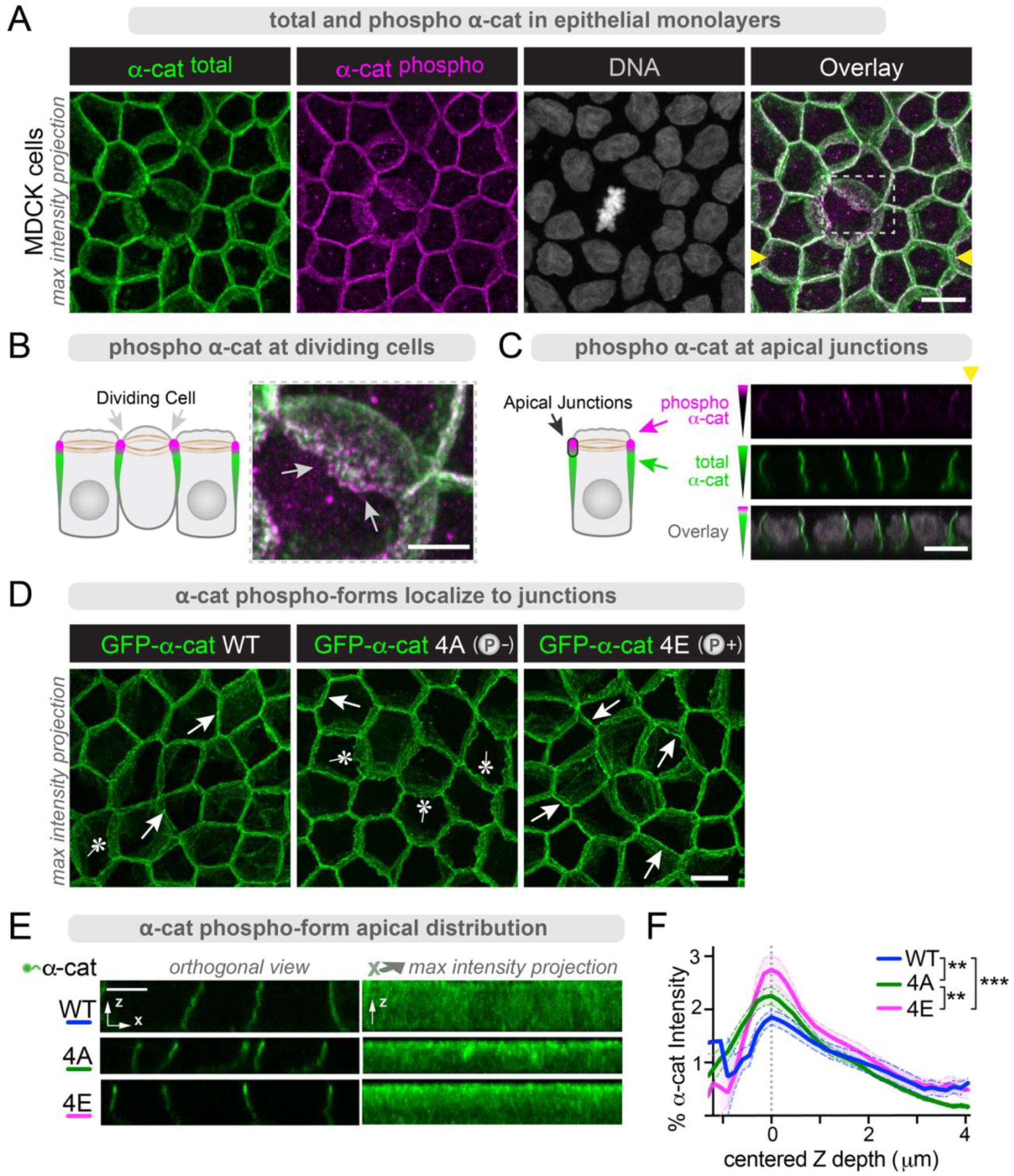
α-cat phosphorylation directs its zAJ enrichment. (**A-C**) Phospho-α-cat localizes to the apical-most portion of adherens junctions. **(A)** Confocal image (z-stack maximum intensity projection) of MDCK monolayer (filter grown) fixed and immuno-stained with antibodies to α-cat (green) and α-cat phosphorylated at S655/T658 (magenta). DNA stained with Hoechst (grayscale). Overlay image of total α-cat (green) and α-cat phosphorylated at S655/T658 (magenta). Scale bar = 10μm. Boxed inset over mitotic cell junction expanded in (**B**); region corresponding to orthogonal x-z section (yellow arrowheads) in (**C**). (**B**) Apical-most portion of zAJ reveals enrichment of α-cat pS655/658 (magenta). Scale bar = 5μm. Schematic shown to left of image. (**C**) Confocal x-y section shows pS655/T658 apical junction enrichment relative to total α-cat (green). Schematic shown to left. Scale bar = 10μm. (**D-E**) α-cat 4E phosphomimic shows intrinsic capacity for zAJ enrichment. (**D**) Confocal images of filter-matured MDCK a-cat KO cells restored with GFP-α-cat phospho-forms. Arrows show apical enrichment of GFP-α-cat signal; asterisks show less enrichment. Scale bar = 10μm. (**E**) Orthogonal views of monolayers show differences in individual junction distribution, captured across whole FOVs with maximum intensity projection, quantified in (**F**). α-cat 4E has the highest standardized intensity value at apical junctions. α-cat 4A is less able to apically pack, showing broader, less condensed signal intensity. n=11 FOV. **P<0.01, ***P<0.001 by multiple t-tests and ***P<0.001 by ANOVA.

**Figure 2:**
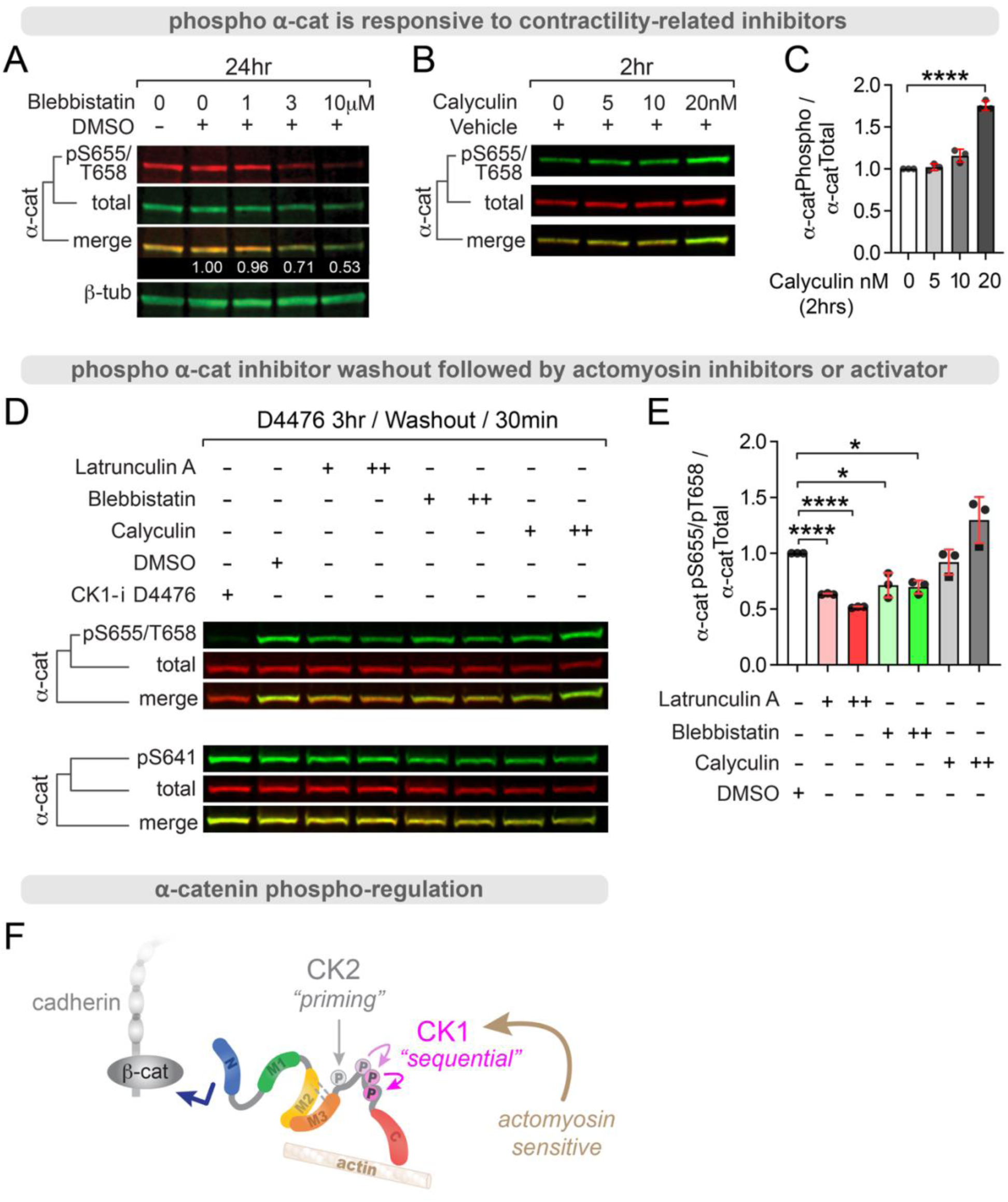
α-cat phosphorylation is actomyosin sensitive. MDCK cells treated with inhibitors (A) and activators (B) of actomyosin contractility show changes to α-cat CK1 phospho-sites pS655/pT658 in total cell lysates. (**A**) Myosin ATPase inhibitor, blebbistatin, dose-dependently inhibits α-cat pS655/pT658 by immunoblot analysis. Numbers (white, merge image) reflect pS655/T658 signal (red) intensity relative to total α-cat intensity (green). β-Tubulin is used as housekeeping protein control. (**B**) Representative immunoblot showing actomyosin contractility promoter and phosphatase inhibitor Calyculin A enhances these same phosphorylations. (**C**) Graph shows quantification of bioreplicate immunoblots. ****P<0.0001 by t-test. (**D**) Actomyosin regulators impact acute recovery of α-cat phosphorylation after CK1 inhibition. MDCK treated with the CK1 inhibitor (D4476) for 3hrs, followed by wash-out (30-minutes) in DMSO, latrunculin, blebbistatin and Calyculin. Representative immunoblot shows that actin inhibitors latrunculin and blebbistatin attenuate recovery of pS655/T658, while Calyculin enhances recovery. (**E**) Graph quantification of bioreplicates. *P<0.05, ****P<0.0001 by multiple t-tests (**F**) Schematic of dual-kinase phosphorylation scheme, where CK1 sites are actomyosin sensitive.

### α-cat phospho-mimic shows enhanced zAJ enrichment and M-domain accessibility

To establish whether phospho-α-cat enrichment at zAJs is causal to or a consequence of its modification by apically localized enzymes, we generated GFP-α-cat fusions that replicate phospho-mutant (4A), phospho-mimic (4E), and wild-type (WT)-α-cat capable of dynamic phosphorylation. These constructs were introduced into canine-α-cat CRISPR-KO MDCK cells via lentiviral delivery. Expression levels were ensured by flow cytometry sorting cells with similar GFP fluorescence intensities and validation by immunoblotting (Fig. S1). We matured epithelial monolayers on high density pore polycarbonate filters and processed for immunofluorescence after barrier maintenance for ∼2 weeks post-seeding. This ensures junction phenotypes are not simply due to differences in monolayer establishment rates after trypsinization. Using confocal-imaging analysis, we noticed that α-cat KO MDCK cells restored with phosphomimic α-cat (GFP-α-cat 4E) showed stronger apical enrichment (arrows) compared with phospho-mutant α-cat (GFP-α-cat 4A), which generally lacked apical enrichment (asterisks) (Fig. 1D). To address robustness and reproducibility of this phenotype, we quantified α-cat apical enrichment across entire monolayers (Fig. 1E). Plotting α-cat mean fluorescence intensity by z-depth (Fig. 1F), we found that the α-cat 4E mutant displayed significantly greater apical enrichment than α-cat 4A or WT α-cat. These data suggest α-cat phosphorylation is not merely a consequence of zAJ localization, but is intrinsically required for this localization.

Actomyosin tension-dependent α-cat M-domain opening is considered required for zAJ enrichment^45^, but whether phosphorylation of the P-linker can prime α-cat for force-dependent opening has remained untested. We sought to address this question using an established *in vitro* kinase assay for α-cat dual phosphorylation by CK2 and CK1^66^. When candidate P-linker sites available for modification are mutated to alanine residues (α-cat P-mutant), radioactive ^32^[P]-incorporation by CK2/CK1 incubation is completely blocked (Fig. 3A), as previously shown^66^. Curiously, a P-linker phospho-mimic mutant showed residual, but significantly enhanced ^32^[P]-incorporation compared with the α-cat phospho-mutant, raising the possibility of phosphorylation at other sites in α-cat (Fig. 3A). While α-cat P-linker sites are the most abundant phospho-sites detected in α-cat^60–66^, we and others found evidence of lower abundance phosphorylations within the M-domain region (pS453/455, pS507 and pS634) (Fig. S2). When we compared ^32^[P]-incorporation of the α-cat P-linker phospho-mimic to an α-cat mutant that cannot be phosphorylated at both P-linker and M-domain sites (α-cat Phospho-null), only background levels of radioactivity were detected (Fig. 3B). These data reveal that phosphorylation of α-cat’s P-linker region can favor accessibility of its M-domain to modification by kinases (Fig. 3C). This increased M-domain accessibility suggests that charge modification of the P-linker may allosterically prepare α-cat for activation and enhanced zAJ enrichment in cells.

**Figure 3:**
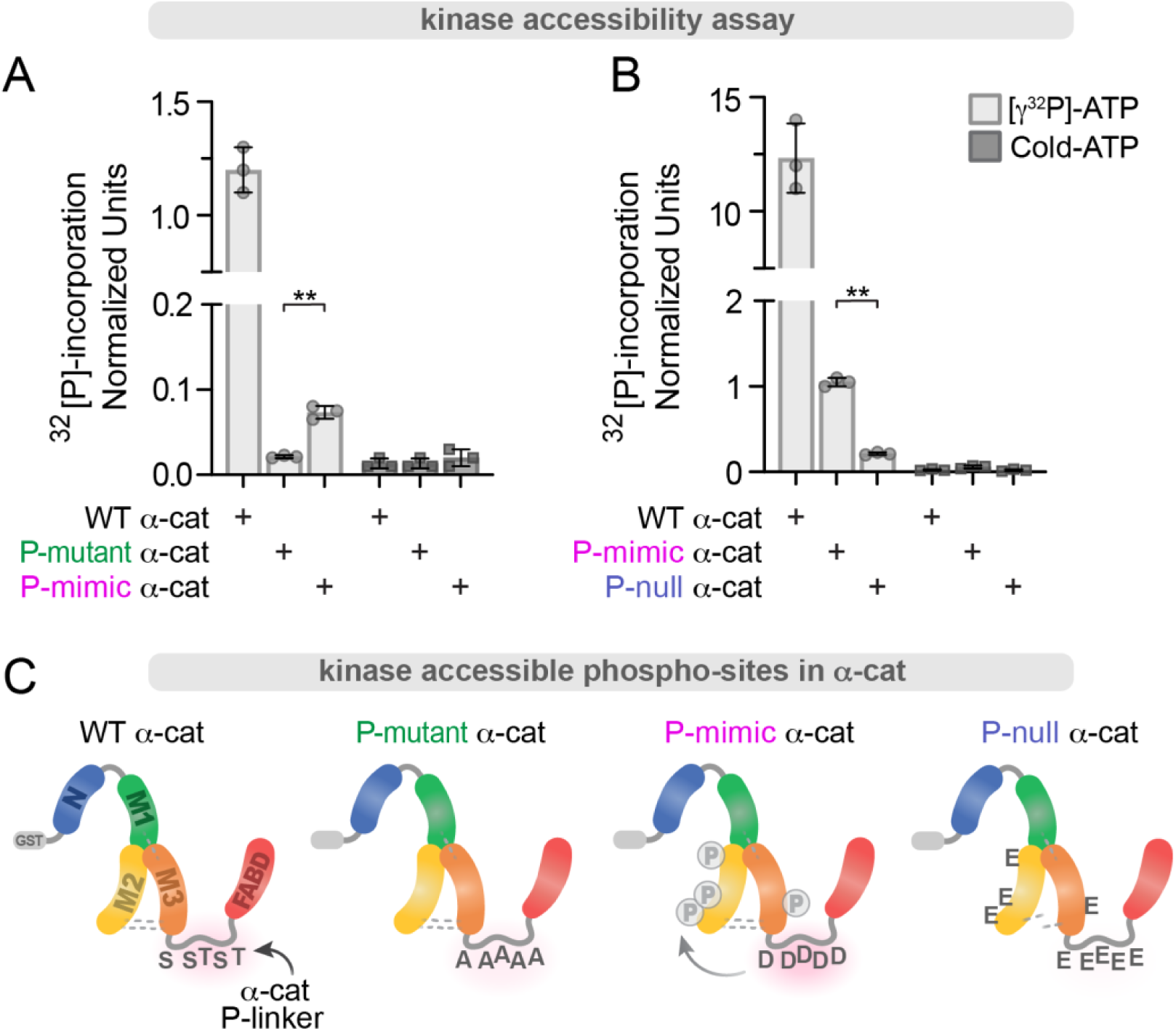
α-cat P-linker phospho-mimic promotes M-domain accessibility. (**A-B**) Graphs show quantification of γ32[P]-incorporation after incubation of α-cat forms (**C**) with CK2, CK1 and γ32[P]-ATP. (**A**) Residues in the P-linker are responsible for the majority of α-cat phosphorylations *in vitro* (comparison of γ32[P]-incorporation between WT versus P-mutant (green) or P-mimic α-cat (magenta)). Despite the same S/T residues in the P-linker being mutagenized, the P-mimic form shows increased phospho-signal compared to P-mutant α-cat. (**B**) Graph quantification of γ32[P]-incorporation of WT α-cat compared to P-mimic α-cat (magenta), and a P-null mutant (blue), where pS/pT residues previously mapped in the M-domain were also removed. Evidence that the P-null α-cat eliminates residual phospho-signal of the P-mimic α-cat, suggests the difference in P-mimic and P-mutant signal (**A**) is due to modest allosteric activation of the M-domain. Unlabeled ‘cold’ ATP was used as a negative control across the time course (panel B). **P<0.01 by t-tests.

### α-cat phospho-mimic enhances Afadin recruitment

Afadin is an established zAJ constituent^3^, α-cat-binding partner^15,43^, and recently predicted to engage α-cat’s M3-domain through a short helical portion of Afadin’s intrinsically disordered region^43,58^. Since the P-linker region of α-cat lies just C-terminal to the M3-domain, we speculated that α-cat phosphorylation might impact its co-localization with and/or binding to Afadin. We used confocal microscopy to image mature MDCK monolayers restored with WT, 4A (mutant) and 4E (mimic) α-catenins. We also included two mutants that interrupt salt-bridge interactions (KRR>AAA, and orthogonal, DDD>AAA mutants), leading to constitutive opening of the M-domain and hyper-recruitment of α-cat binding partners^33–35,71–73^. While all α-cat constructs generally colocalize with Afadin at the zAJ (Fig. 4A), only WT and 4E α-catenins show full co-localization with Afadin at the apical-most zAJ (Fig. 4B-C). In contrast, the α-cat 4A mutant shows only 75% signal overlap at zAJ, suggesting less co-localization with Afadin. Although α-cat salt-bridge mutants show somewhat less co-localization with Afadin at zAJs, this is likely because Afadin and α-cat KRR and DDD mutants are hyper-enriched at tricellular junctions (Fig. 4A, arrows), which are known to extend deeper along the lateral membrane. We sought to quantify differences in Afadin recruitment further using a simple junctional enrichment measurement, where fluorescence intensity from 1μm circular ROIs were measured from bicellular junctions (BCJ) and the adjacent cytoplasm (Cyto). Although differences are modest, this approach confirms that the α-cat 4E mutant shows significantly greater enrichment of Afadin to zAJs than WT or 4A α-cat (Fig. 4D). α-cat KRR and DDD salt-bridge mutants also show greater Afadin recruitment than WT, consistent with their persistently open M-domain structure. These data suggest that α-cat phosphorylation is required for robust colocalization with its zAJ partner, Afadin.

**Figure 4:**
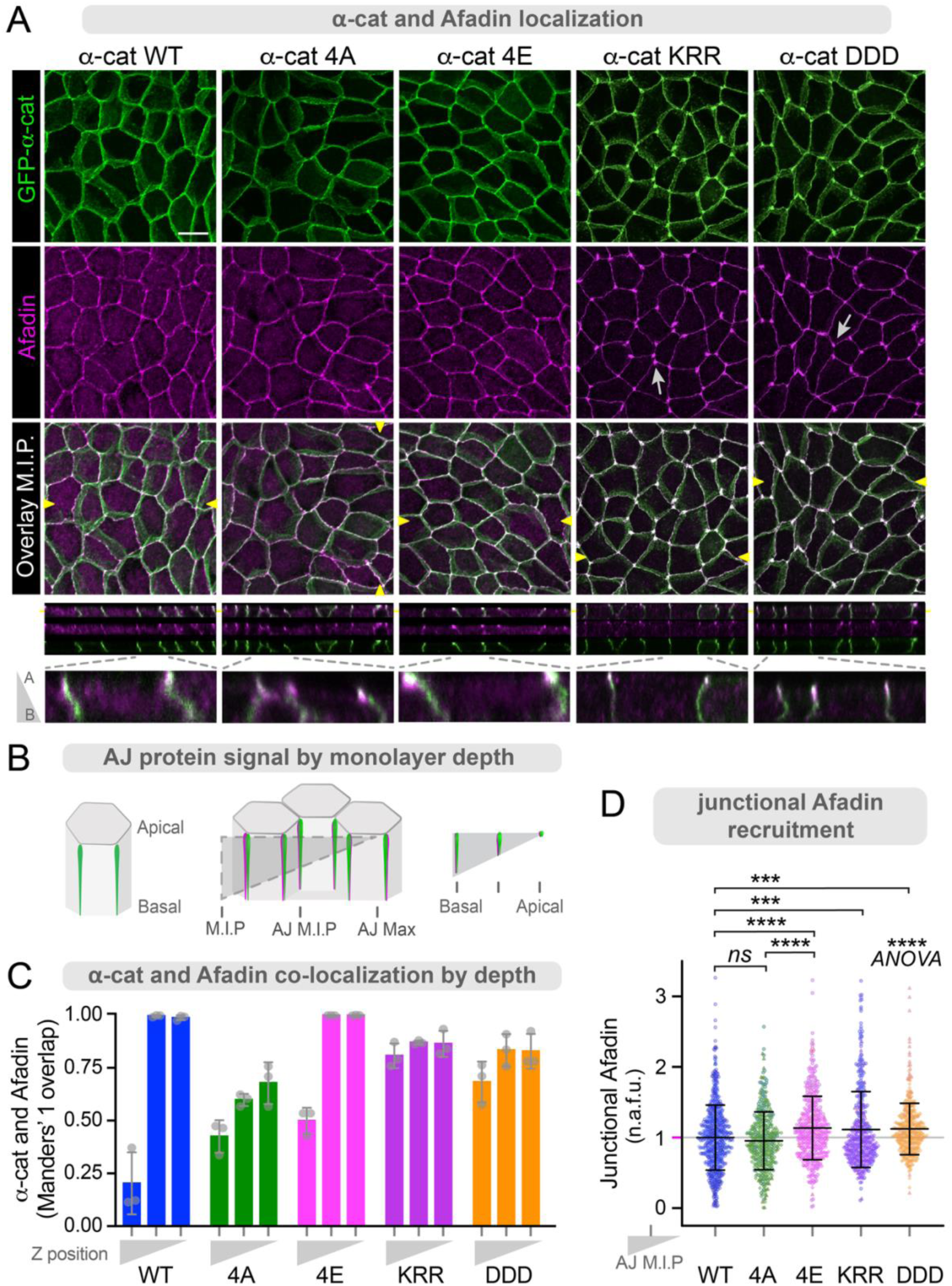
α-cat P-linker phospho-mimic promotes zAJ recruitment of Afadin. (**A**) Confocal images of MDCK monolayers with restored GFP-α-cat forms fixed and immuno-stained for Afadin (magenta). *En face* images are maximum intensity projections (M.I.P.). Scale bar = 10μm. Corresponding orthogonal planes (yellow arrowheads) are shown with insets zooming in on individual junctions. (**B**) Schematic shows cell/image depth for α-cat/Afadin colocalization analysis. Confocal Z-stacks were analyzed as M.I.P.s to include basal signal, AJ M.I.P.s to isolate junction signal, and single slices focused on maximum apical α-cat signal. (**C**) Manders’ co-localization by monolayer Z-depth. The fraction of α-cat signal that overlaps with Afadin signal is highest in apical portions of WT and 4E monolayers. (**D**) Junctional Afadin enrichment was measured using 1μm circular ROIs taken from bicellular junctions, subtracting adjacent cytoplasm signal. Normalized fluorescence was plotted, with symbols corresponding to junctions from 3 different experiments with 6+ FOV each. ***P<0.001, ****P<0.0001 by t-tests. ****P<0.0001 by ANOVA.

### α-cat M3-domain point mutants inhibit Afadin-binding in vitro

To better understand the mechanosensitive interaction between Afadin and α-cat, we sought to identify the minimal Afadin-binding region on α-cat. Since the α-cat-Afadin interaction is favored by the salt-bridge disrupting R551A mutation or mutants that release the M-domain from auto-inhibition^15^, we posited that the α-cat M-domain contains an Afadin-binding site. We tested this idea by performing pulldown assays using GST-Afadin-<u>C</u>oiled-<u>c</u>oil <u>α</u>-cat <u>b</u>inding region (Afadin-CC-ABR; amino acids 1391-1449), based on larger regions previously shown to interact with α-cat (Marou et al^43^, amino acids 1378-1473); Sakakibara et al^15^, amino acids 1393-1602), incubated with various α-cat M-domain constructs (Fig. 5A-B). For this initial mapping, we used the neural version of α-cat (*CTNNA2*, or αN-cat), where the salt-bridge disrupting R551 equivalent mutation is R549A (Fig. 5A). While the α-cat-M1-3-WT interaction with GST-Afadin-CC-ABR was barely detectable by Coomassie staining, the α-cat-M1-3-R549A showed markedly increased interaction with GST-Afadin-CC-ABR (Fig. 5B). These data confirm that the M-domain of αN-cat, like αE-cat, must be opened to bind Afadin. These data also suggest that the Afadin-binding site is located within the M-domain, where its accessibility to Afadin is regulated by the same open-closed conformational dynamics of the α-cat-M domain that controls vinculin-binding to the α-cat-M1 region^71,72^. Accordingly, use of WT and R549A variants of α-cat-M2-3 (where the M1 deletion effectively destabilizes the closed M-domain conformation), displayed uniformly strong binding to GST-Afadin-CC-ABR. Since the αN-cat-M3 alone binds equivalently to GST-Afadin-CC-ABR, we conclude that α-cat-M3 constitutes the core Afadin-binding region.

**Figure 5:**
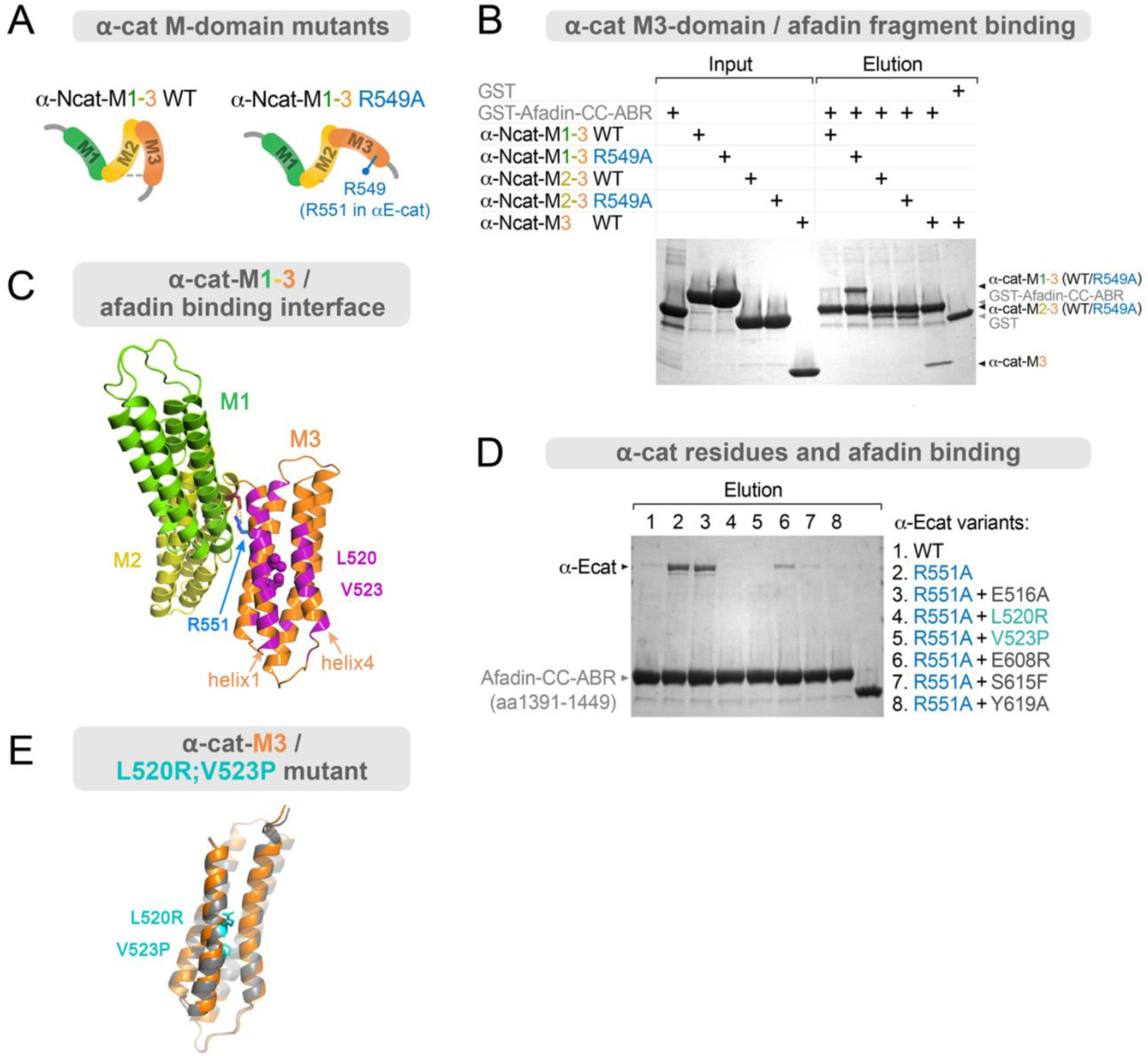
α-cat-M3 interacts with Afadin coiled-coil domain residues 1391-1449. (**A**) Schematic of truncated α-cat forms used for Afadin interaction mapping. αN-cat was used for domain mapping in A-B. The commonly used αE-cat M-domain salt-bridge disrupting mutant R551A corresponds to R549A in αN-cat. (**B**) GST-Afadin-<u>α</u>-cat-Binding-<u>R</u>egion (Afdn-CC-ABR, residues 1391-1449) pulldown assay with αN-cat M-domain fragments (prey proteins). Various M-fragments (M1-3, M2-3 and M3) of αN-cat were mixed with GST control or GST-Afdn-CC-ABR pre-bound to glutathione resin, followed by washing and elution with free-glutathione. Comparison of αN-cat M1-3 WT and αN-cat M1-3 R549A co-sedimenting with GST-Afdn-CC-ABR indicates that the αN-cat Afadin-binding site is autoinhibited in αN-cat WT, whereas it becomes more accessible to Afdn-CC-ABR in the R549A variant or other M-fragments missing the M1-domain. (**C**) Afadin-binding site of αEcat (indicated by magenta) determined by NMR titration experiments (Fig. S4A) is mapped on the surface of M3-domain, containing L520 and V523. The R551A mutation is expected to break the central salt bridge to release M3 from an autoinhibited/M1-proximal state to increase Afadin binding. (**D**) GST-Afdn-CC-ABR pulldown assay with various full length αE-cat variants. αE-cat R551A clearly binds to GST-Afdn-CC-ABR, whereas only a negligible amount of bound αE-cat WT was detected. Afadin-binding site mutations (E516R, L520R, V523P, E608R, S615F and Y619A) were individually combined with R551A to examine the effects of putative mutations under increased afadin binding. L520R and V523P mutations negated Afadin binding. (**E**) Alphafold predictions of αE-cat structure with L520R and V523P point mutations, showing that these mutations do not perturb the overall αE-cat M3 structure. αE-cat M3 folds into helical bundles, with WT (orange) and mutant (WT-ΔAfdn, gray) models overlaid for comparison.

We next sought to define the α-cat-M3-Afadin interface in the context of a full-length M-domain activated αE-cat, leveraging the R551A salt-bridge perturbing mutant to constitutively open α-cat for in-solution binding to Afadin (Fig. 5C). Guided by NMR titration analysis, which revealed residues altered by α-Ncat-M3-Afadin binding (Fig. S3A-B), we mutated six individual α-cat residues corresponding to the first and fourth helices of α-cat M3 and performed *in vitro* pulldown assays with GST-Afadin-CC-ABR (Fig. 5D). As expected, the R551A mutation leads to a clear increase in the binding of αE-cat with GST-Afadin-CC-ABR compared to WT α-cat. When we combine the R551A mutation with an additional Ala substitution at the Afadin-binding site, five out of six mutations markedly reduced the αE-cat-Afadin interaction (Fig. 5D). Given the ability of L520R or V523P mutants to completely block Afadin-binding in dilute solution, and their close proximity, we generated an L520R/V523P double point mutant (Fig. 5E) to interrogate consequences for Afadin-recruitment, zAJ structure and function in cells.

### α-cat M3-helix1 contributes to Afadin recruitment and normal epithelial barrier morphology

To address the consequence of losing this α-cat-M3/Afadin-CC-ABR interaction for epithelial monolayer organization, we restored α-cat KO MDCK cells with an L520R/V523P mutant form of α-cat, hereafter referred to as GFP-α-cat Δ-Afadin (α-cat Δ-Afdn). Although α-cat WT, α-cat Δ-Afdn mutant and endogenous Afadin proteins were similarly well-expressed, we could not validate the contribution of α-cat M3 residues L520/V523 to Afadin-binding by co-immunopreciptation from cell lysates, as negligible levels of Afadin co-associated with either α-cat (Fig. S4), likely because detergent-extractable α-cat is largely auto-inhibited. Thus, as is common for α-cat M-domain effector studies, we turned to quantitative immunofluorescence microscopy. Confocal imaging of mature epithelial monolayers revealed that α-cat Δ-Afdn mutant cells show recessed multi-vertex junctions, which appear as lateral gaps (Fig. 6A, apical x-y plane, yellow arrowheads; x-z gaps, yellow asterisks). These multi-vertex defects were substantially more numerous in α-cat Δ-Afdn mutant compared with WT α-cat monolayers (Fig. 6B). While the α-cat Δ-Afdn mutant does not completely block Afadin recruitment to zAJs, unsurprising since Afadin can interact with other TJ/AJ constituents^11,12,74,75^, bi-cellular junction enrichment was significantly reduced (Fig. 6C-D). These data suggest that the α-cat M3/Afadin-CC-ABR interaction contributes to the balanced distribution of Afadin along zAJs, and is particularly critical for multi-vertex junction organization.

**Figure 6:**
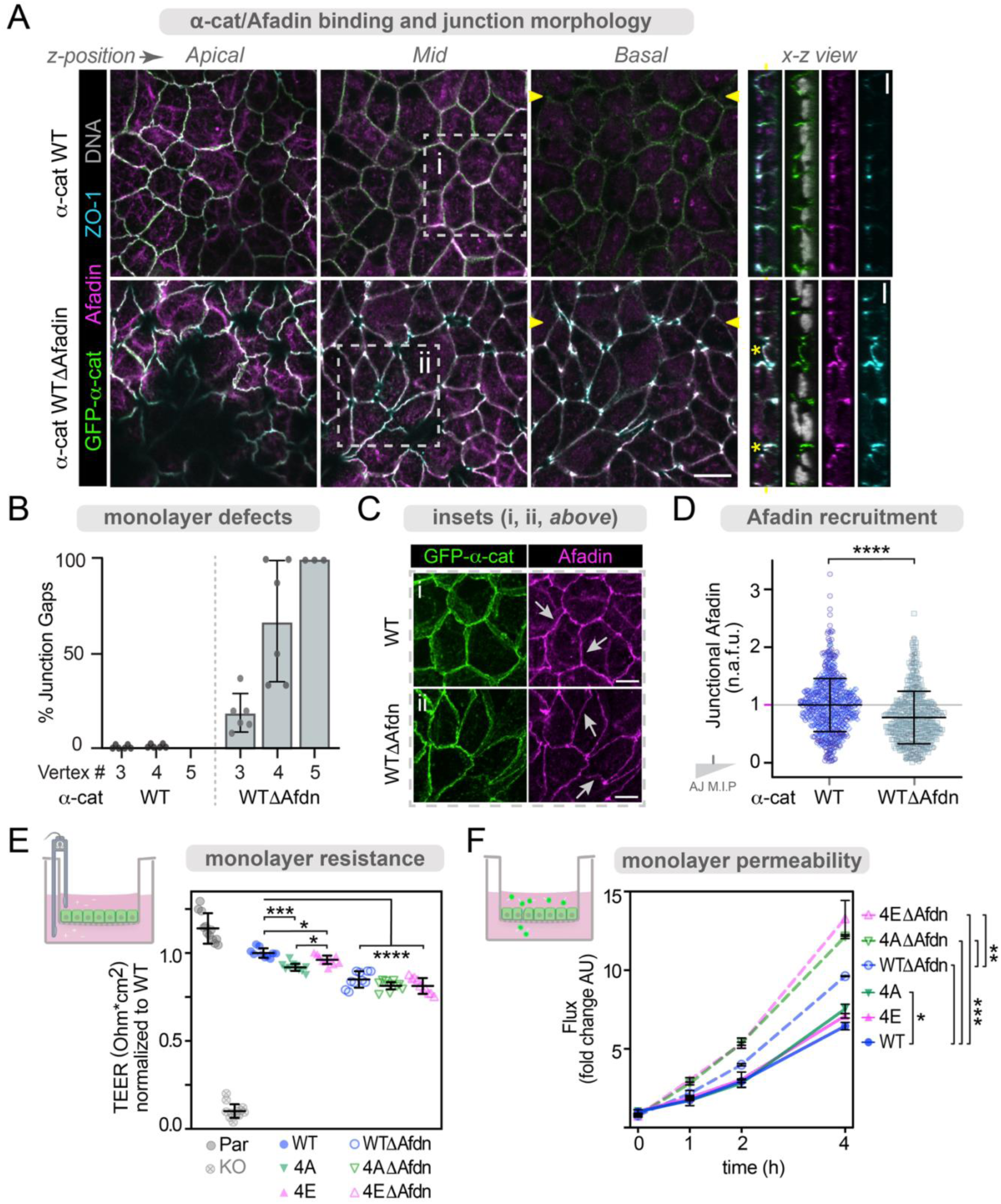
α-cat M3-Δ-Afadin binding mutant interferes with epithelial barrier function. (**A**) Single confocal sections of mature MDCK monolayers reconstituted with GFP-WT α-cat or GFP-WTΔAfadin (L520R V523P) α-cat and immunostained with antibodies to Afadin and ZO-1. Apical junction components Afadin, ZO-1 and α-cat localize to cell-cell contacts in both WT and ΔAfdn monolayers (white signal overlay). α-cat ΔAfdn monolayers have irregular junction signal and dysregulated junction positions, with Afadin and tight junction protein ZO-1 extending basally. Scale bar = 25 μm. Corresponding orthogonal planes (yellow arrowheads) are shown to right, with GFP-α-cat, Afadin and ZO-1 labeling shown separately. Asterisks (yellow) show apical multicellular junction defects, often extending to the basal most aspect of the monolayer. While single apical slices show gaps at junctions, these areas come together as more basal z-planes come into view. Scale bar = 10 μm. Insets zoom in on individual junctions (i and ii) shown in C. (**B**) Percent of junctions with gaps by junction number (e.g. 3 = three-cell-junctions). 0% of 3- or 4-cell junctions in α-cat WT monolayers showed holes in apical confocal z-slices. Multi-cellular junctions in α-cat ΔAfadin monolayers show more junction gaps with increasing cell number. Values plotted are mean +/− SD from 6 FOV. (**C**) Insets i and ii from A, show single color maximum intensity projection views (α-cat, green; Afadin, magenta) of cell-cell junctions. (**D**) Junctional Afadin enrichment was measured using 1μm circular ROIs taken from bicellular junctions, subtracting adjacent cytoplasm signal. Normalized fluorescence was plotted, with symbols corresponding to 3 different experiments with 6+ FOV each. ****P<0.0001 by t-tests. (**E-F**) α-cat phospho- and Afadin-binding mutants show barrier dysfunction, with α-cat P-mimic unable to overcome removal of Afadin-binding. (**E**) α-cat phospho-state and ability to bind Afadin independently contribute to barrier dysfunction by day 7 TEER analysis using a standard volt-ohm meter (schematic, left). Normalized resistance is plotted as mean +/− SD). n=8-10 from 2 biological replicates. *P<0.05, ***P<0.001; ****P<0.0001 by multiple t-tests, and ***P<0.001 by ANOVA. (**F**) Barrier to small molecules detected by FITC-4kDa-Dextran leak (‘Flux’) across day 10 monolayers (schematic, left). FITC leak was measured from the basal side of monolayers over time, with averaged fluorescence from 3 monolayers plotted as mean +/− SD.

A number of groups have generated Afadin KO cell lines^15,16^, where epithelial monolayer phenotypes show little evidence of multi-vertex junctional gaps (Fig. 6A). This raised the possibility that culture conditions could account for differences between Afadin KO and α-cat Δ-Afdn monolayer phenotypes. Alternatively, the α-cat Δ-Afdn mutant might perturb more than the α-cat M3/Afadin-CC-ABR interaction. To address this question, we compared junction phenotypes of Afadin KO MDCK epithelial monolayers matured on glass (Fig. S5) versus filters (Fig. S6). Similar to others^15,16^, we found no evidence of intercellular gaps for either plating condition, although Afadin KO MDCK matured on filters developed an intraepithelial lumen phenotype (Fig. S6B). Of interest, the multi-vertex gap phenotype is only seen when α-cat Δ-Afdn mutant cells are matured on filters (Fig. S6A, arrows), but not glass (Fig. S5A). Afadin shows reduced co-localization with α-cat Δ-Afdn compared with α-cat WT under either plating condition (Fig. S5D vs Fig. 6D). Thus, the multi-vertex gap phenotype appears to be a feature of α-cat Δ-Afdn mutant cell monolayers fully matured on filters. To address the possibility that our α-cat Δ-Afdn mutant might perturb more than the α-cat M3/Afadin-CC-ABR interaction, we modeled this interaction using AlphaFold, particularly since our α-cat Δ-Afdn mutant incorporates a proline residue into the first helix of M3 (L520R/V523P). While the α-cat M3-L520R/V523P mutants can break the helical structure of α-cat M3 helix 1 in isolation, the impact of these point mutations on the larger structure is minimized, suggesting reduced impact on the full-length protein (Fig. S3C-E). Together, these data demonstrate that the α-cat Δ-Afdn mutant interferes with zAJ organization, in part, through Afadin-binding. Since α-cat Δ-Afdn and Afadin-KO phenotypes are distinct, the α-cat Δ-Afdn mutant may impact more than Afadin recruitment.

### α-cat phosphorylation and Afadin-binding are required for epithelial barrier function

While the α-cat 4A mutant shows reduced zAJ enrichment (Fig. 1), M-domain accessibility (Fig. 3) and Afadin co-localization (Fig. 4), whereas the α-cat 4E mutant shows enhanced zAJ enrichment, M-domain accessibility and Afadin recruitment, it remained unclear whether these modest differences are physiologically meaningful. Towards this end, we compared epithelial barrier characteristics across all of our expression-matched cell lines. We found that α-cat 4A monolayers show significantly less epithelial resistance than WT or α-cat 4E cells (Fig. 6E), along with a modest increase in small molecule flux (Fig. 6F), consistent with the morphological differences described above. Curiously, while α-cat 4E monolayers show higher epithelial resistance than α-cat 4A cells, the phospho-mimic shows less barrier resistance than WT cells (Fig. 6E), despite its enhanced zAJ enrichment and Afadin recruitment (Figs. 1, 4). Thus, while α-cat 4A and α-cat 4E mutants display distinct junction phenotypes and Afadin recruitment, an α-cat capable of dynamic on-off phosphorylation may be most optimal for epithelial barrier function.

We also compared α-cat 4A and 4E barrier phenotypes against the α-cat Δ-Afdn mutants (Fig. S1, immunoblot). Interestingly, the modestly increased epithelial resistance seen in α-cat 4E versus α-cat 4A monolayers was not observed between α-cat 4A Δ-Afdn and α-cat 4E Δ-Afdn “double-mutant” monolayers (Fig. 6E). These data suggest that while the improved functionality of α-cat 4E over 4A may be via Afadin, it is also clear that all α-cat Δ-Afdn mutants show more greatly attenuated epithelial barrier resistance, underscoring the importance of α-cat’s M3-helix 1 to barrier function. Additional evidence that α-cat 4A Δ-Afdn and α-cat 4E Δ-Afdn “double-mutant” monolayers show significantly greater permeability than the α-cat Δ-Afdn alone (Fig. 6F), suggests the possibility that α-cat P-linker and Afadin-binding-regions may also independently contribute to epithelial barrier function (see Discussion).

### Loss of α-cat phosphorylation impairs development and leads to hydrocephalus

To investigate the role of α-cat P-linker phospho-sites (S641, S652, S655, and T658) in mammalian tissue development, we generated mice incapable of phosphorylation at these residues using site-specific CRISPR mutagenesis ((*Ctnna1*^4A/+^; Fig. 7A; Fig. S7). Two independent founder lines were intercrossed to generate *Ctnna1*^4A/4A^ homozygous knock-in mutant mice. Both lines underrepresented the *Ctnna1*^4A/4A^ homozygous genotype at the time of weaning, in part, due to post-natal demise (Fig. 7B). While *Ctnna1*^4A/4A^ mice were significantly smaller than WT littermates (Fig. 7C), no obvious differences in heart, lung, liver, kidney, or gut were detectable. Since *Ctnna1*^4A/4A^ mice failed to reach adulthood (10-weeks; 0/22 *Ctnna1*^4A/4A^ weanlings) and needed to be humanely euthanized due to hydrocephalus (Fig. 7D), we sought to quantify this phenotype by histology. Using Nissl-stain to ensure comparison of ventricle sizes from matched brain depths (Fig. 7E), we demonstrated that *Ctnna1*^4A/4A^ ventricles are significantly larger than *Ctnna1*^WT/WT^ mice (Fig. 7F). These data show that α-cat phosphorylation contributes to normal post-natal development and is particularly critical for establishing and maintaining ventricles of the brain.

**Figure 7:**
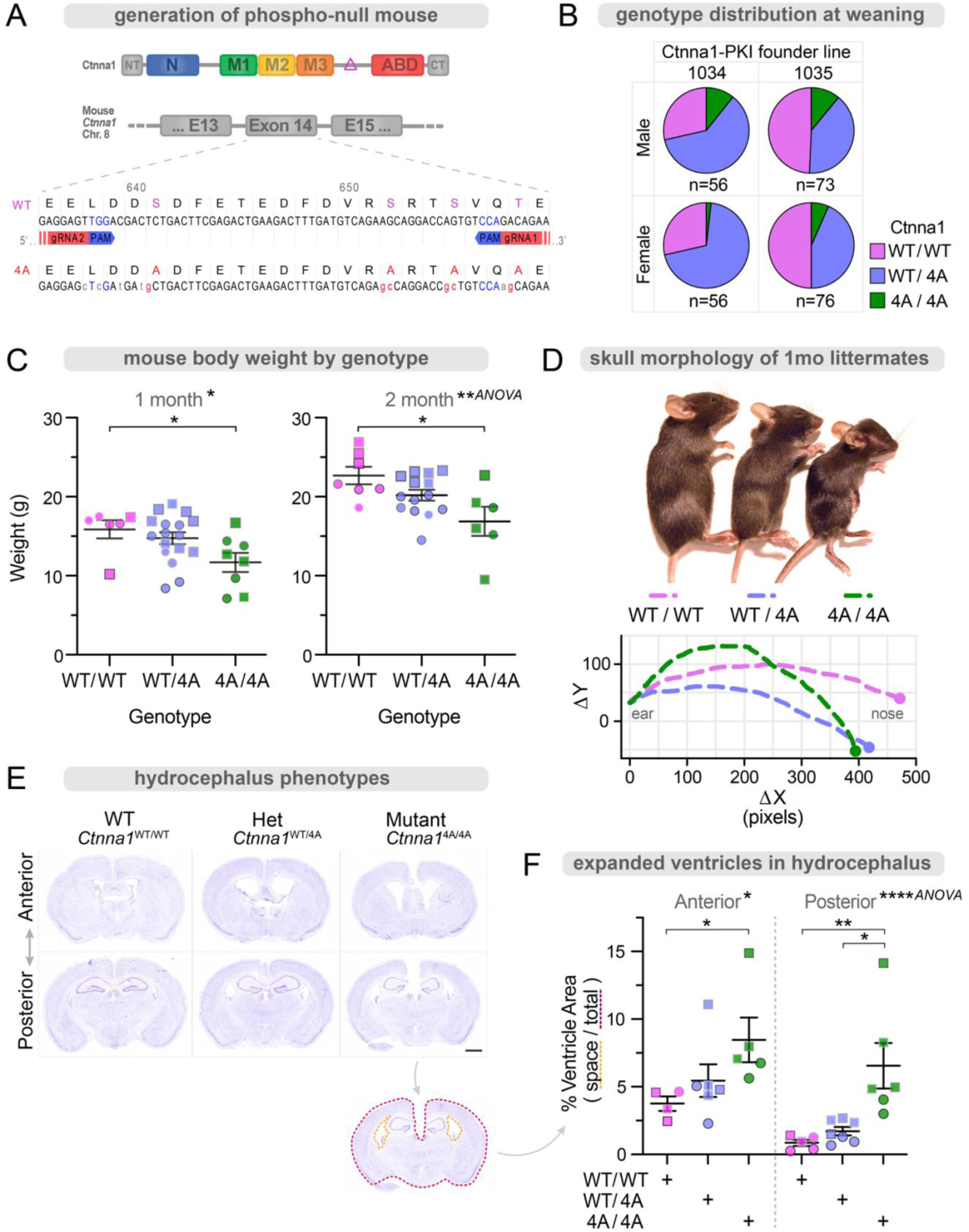
Targeted loss of α-cat phosphorylation impairs development and leads to hydrocephalus. (**A**) Generation of α-cat phospho-mutant mouse. A CRISPR-Cas9 system was used to remove the linker phospho-sites from germline αE-catenin (*CTNNA1*). Two sets of guides were used in two different founders to eliminate the possibility of off-target effect. Both lines were backcrossed 9+ times, and specific α-cat phospho-site substitutions were validated at the genetic, transcriptional, and protein level. Heterozygous P-mutant carriers were crossed to create homozygous P-mutant mice. Data represent a shared lethal mutant phenotype of both founder lines. (**B**) Mice from het x het crosses were genotyped at the time of weaning. Mutant (4A/4A) mice were observed less frequently than Mendelian genetics would predict, indicating some level of lethality at the embryonic or post-natal stage. n=261. (**C**) Mutant mice showed decreased overall weight. Genotyped littermates were weighed before experimental use, with different symbol outlines indicating different litters. Squares indicate males; circles indicate females. Plot shows mean +/− SEM. *P<0.05 by t-test. **P<0.01 by ANOVA. (**D**) Mutant mice show impaired development, often accompanied by morphologic changes to the head and skull consistent with hydrocephalus, requiring humane sac and leading us to investigate this early cause of demise. (**E**) Brains from mutant mice have increased ventricular space, consistent with longstanding hydrocephalus. Perfused brains were excised, fixed, embedded, sectioned, and Nissl stained. Stereotypic stain patterns were used to identify matched anterior and posterior tissue sections across samples. Scale bar = 1mm. (**F**) Ventricular area and total brain area were measured, with an increased ratio of free ventricular space representing hydrocephalus. Different symbol outlines indicate different litters. Squares indicate males; circles indicate females. Plot shows mean +/− SEM. *P<0.05, **P<0.01 by t-test. *P<0.5, ****P<0.0001 by ANOVA.

## DISCUSSION

Historic electron micrographs reveal adherens junctions as highly regular and compact structures^2^, but molecular rules and interactions for how AJ are progressively built remains incompletely defined. Focusing on the central mechanosensitive component of AJ, α-cat, and one of its effectors, Afadin, we provide evidence that α-cat integrates mechanical and chemical signals to promote zAJ organization and function. We previously identified an evolutionarily-conserved phosphorylation relay in a flexible region of α_cat_ linking its tension-sensitive F-actin and Middle-domains, where these sites are required for strong cell-cell adhesion in mammalian cells and fly development^66^, but mechanistic understanding remained elusive. Using phospho-specific antibodies, we find that phospho-α_cat_ enriches at the apical-most portion of adherens junctions, a structure proximally aligned with tight junctions. Evidence this apical enrichment can be phenocopied by a phospho-mimic α-cat, along with evidence these phosphorylations are reduced by inhibitors of actomyosin contractility, suggest α_cat_ phosphorylations are tension-dependent and intrinsically drive α_cat_ junctional enrichment. Further *in vitro* evidence that phospho-mimic, but not phospho-mutant α_cat_, adopts a more open Middle-domain, suggests α-cat phosphorylation may drive junction organization through recruitment of a key binding partner. We show that Afadin is one such binding partner, identifying residues along α-cat M3 helix 1 and 4 that perturb Afadin binding in solution. Using an α-cat M3-helix 1 double-mutant, we show that Afadin recruitment is reduced at bicellular zAJs and greatly perturbs multi-cellular junctions. Since junction maturation and zAJ separation occurs over time^3^, we suggest a stepwise model where α_cat_ integrates mechanical and biochemical signals to progressively drive its epithelial apical junction enrichment, alignment and function (Model Figure).

We do not yet understand the endogenous apical actomyosin contractility pathways that promote α-cat P-linker phosphorylation. Numerous scaffold regulators of Rho-ROCK and myosin control AJ contractility (e.g., Shroom, LUZP, cingulin, Lmo7, *Ccdc88c*/Daple), where all of these proteins are found in proximity proteomic screens of the cadherin, α-cat or Afadin^76–78^. The degree to which these scaffold regulators collectively or selectively regulate α-cat phosphorylation will require further investigation.

There are a few ways to view how α-cat phosphorylation promotes zAJ enrichment. Previous evidence that an α-cat P-linker phospho-mimic alters the tryptic peptide pattern compared to wild-type or phospho-mutant α-cat, effectively preserving the linker-region^66^, suggests that phosphorylation may promote local structure in a previously unstructured region. Since the P-linker joins mechanosensitive M-domain and actin binding-domains, introducing structure could reduce flexibility between these regions. This could in turn favor close packing of α-cat adhesive complexes at dense zAJ, as seen in historical electron micrographs^2^.

Curiously, we find that α-cat P-linker phosphorylation is not only downstream of actomyosin contractility, but favors accessibility of its M-domain to modification by kinases. Regarding the former, we previously showed that α-cat is subject to a dual-kinase phosphorylation mechanism, where α-cat is first modified at S641 by CK2, followed by CK1-dependent modification of residues S652, S655 and T658^66^. Evidence presented here, where this latter triplet of phosphorylations is actomyosin sensitive, appears consistent with previous evidence that these sites are less accessible to modification in full-length α-cat, compared with an α-cat lacking N-terminal through M2-regions^66^. This suggests that α-cat binding to F-actin makes the P-linker more accessible to modification by CK1. Evidence that α-cat P-linker charge modification can also enhance M-domain accessibility raises the possibility that progressive charge modification of the P-linker may allosterically prepare (i.e., prime) α-cat for mechano-activation and enhanced zAJ enrichment in cells. Indeed, we previously showed that α-cat modification by CK1 is sequential, where mutation at S652 blocks further modifications at S655 and T658^66^, but the rationale for a sequentially-dependent modification scheme remained unclear. Our evidence that these phosphorylations are actomyosin sensitive, while also favoring M-domain opening, raises the possibility that these phosphorylations function as a ratchet-like mechanism, with each phosphorylation promoted by increasing force thresholds, which in turn progressively prepares the M-domain for effector-binding.

We chose to focus on Afadin as an α-cat M-domain effector impacted by α-cat phosphorylation for a few reasons. First, Afadin is a multi-domain scaffold and α-cat-binding protein previously shown to promote the accumulation of α_cat_ and F-actin to zAJ^3,15,41,79–82^. While Afadin is perhaps best known for its ability to interact with nectin-family adhesion receptors^83^, recent super-resolution imaging studies show that Afadin-nectins localize to the apical-most portion of zAJ, apical to catenin/cadherin proteins and basal to TJ proteins^3^. This apical zAJ enrichment of Afadin is consistent with what we observe in MDCK cells: Phospho-α-cat is enriched at zAJ, and an α-cat phospho-mutant shows less zAJ enrichment and colocalization with Afadin. Together with evidence that Afadin binds directly to α-cat’s M3 regi on^15,41,43^, which is proximal to the P-linker, we reasoned Afadin may be a phosphorylation-dependent effector ofα-cat critical for epithelial barrier function. We identified residues in the α_cat_ M3-domain required to bind the Afadin-<u>C</u>oiled-<u>c</u>oil <u>α</u>-cat <u>b</u>inding region (Afadin-CC-ABR; amino acids 1391-1449), suggesting that Afadin engages interactions with α-cat M3 helices 1 and 4, consistent with recent modeling of this interaction^58^. Critically, these residues are only available to bind Afadin when the α-cat M-domain is opened, as with salt-bridge disrupting mutants. We validated the contribution of two residues, L520 and V523, in context of a full-length α-cat using a double-mutant approach, showing that this mutant reduces Afadin recruitment to bicellular junctions and perturbs organization at multi-vertex junctions. Curiously, our α-cat Δ-Afdn mutant does not phenocopy the full Afadin KO phenotype. Whether this is due to differing compensatory mechanisms, or that the cat Δ-Afdn L520/V523 mutant also impacts the conformation regulation of α-cat and/or binding of other effectors will require further investigation.

We used standard transepithelial resistance and flux assays to determine functional consequences of the seemingly modest changes in α-cat enrichment at zAJ and co-localization with Afadin. We went into these experiments with a simple prediction: If α-cat P-linker phosphorylation serves as a “switch” for M-domain opening and Afadin recruitment, the α-cat 4A and α-cat Δ-Afdn mutants should lead to similar barrier dysfunction. However, we did not observe this. Instead, we found that the α-cat Δ-Afdn mutant showed greater barrier leak than the α-cat 4A mutant, and that α-cat-P-linker/Δ-Afdn double-mutants showed additive increases in leak. These data reveal that *both* P-linker and Afadin-binding regions are required to seal the barrier, suggesting these regions independently contribute to barrier function. Since this model is not consistent with α-cat P-linker phosphorylation serving as a simple molecular switch for M-domain opening, we considered alternative explanations. In a companion study *(Sagi et al. [Weis, WI, Dunn, AR], in preparation)*, we show that the α-cat P-linker phosphorylation leads to greatly enhanced binding to ZO-1. Since ZO-1 has binding-sites on Afadin that are proximal to where α-cat M3 binds, we favor a model where that α_cat_ phospho-linker coordinates zAJ enrichment, TJ alignment and epithelial barrier function through promoting a trimolecular complex^3,15,41,79–82^.

We previously showed that α_cat_ P-linker phospho-sites are conserved from *Drosophila* to humans and required for adult fly development^66^, but their requirement for mammalian tissue development remained untested. Here, we generated knock-in mice incapable of phosphorylating α-cat at S641, S652, S655 and T658, and found these mice develop a highly penetrant hydrocephalus phenotype. Mutation or loss of a number of apical junction proteins has been linked to congenital hydrocephalus in humans (*L1CAM, MPDZ, CRB2* and *CCDC88C*/Daple)^84–88^. Since we recently identified the latter protein, Daple, as a proximity partner of wild-type more than a salt-bridge disrupting mutant α-cat^77^, and loss of other zAJ proteins or α_cat_ proximity-partners can lead to hydrocephalus in mice ^89–91^, we speculate that *Ctnna1*^4A/4A^ mice develop hydrocephalus through perturbing a particular feature of neuroepithelial zAJ integrity, which forms a critical barrier between brain cortex and fluid-filled ventricles. Evidence that most *Ctnna1*^4A/4A^ organs appear largely normal (e.g., kidney, lung, intestine; not shown) raises the possibility that tissue-specific differences in mechanical load on zAJ may explain the vulnerability of neuroepithelia to loss of α-cat phosphorylation. Importantly, we cannot rule out roles for α-cat phosphorylation in limiting leak across other barrier cell types, such as endothelia^92,93^. Pinpointing the cell-type through which loss of α-cat phosphorylation leads to hydrocephalus will require new genetic tools to selectively prevent α-cat phosphorylation across neuroepithelial versus endothelial cell-types.

In summary, we show that phosphorylation of a mechanosensitive junction scaffold protein both depends on actomyosin contractility, and primes this scaffold for force-dependent opening and effector recruitment, suggesting how junction structures can be built through integrating mechanical and chemical signals.

## STUDY LIMITATIONS

This study relies on in-solution biochemistry and structurally informed mutant strategies to address the contribution of α-cat phosphorylation and Afadin-binding to zAJ organization and function using the MDCK α-cat KO/reconstitution system. While we credential α-cat L520R/V523P as a Δ-Afdn-binding mutant, other amino acids localizing to α-catM3-helix 4 also contribute to binding, suggesting that the α-catM3/Afadin-CC-ABR may benefit from further refinement. Lastly, the seemingly modest impact of α-cat P-linker phosphorylation in the MDCK cell system was subjected to the robust physiological test of mouse development, where we show a clear role during post-natal brain development. While the hydrocephalus phenotype was observed in two independent founder lines, litter sizes were small (4-5 pups). Thus, we cannot rule out a contribution of the *Ctnna1*^4A/4A^ mutant to early embryonic lethality.

## METHODS

### Lentiviral plasmid generation, constructs and reagents

All GFP-α-cat constructs were synthesized by Vectorbuilder using the αE-catenin human *CTNNA1* sequence. The ability of eGFP to spontaneously dimerize and potentially impact α-cat functions was disrupted by incorporating an A206K mutation^94^. Synthesized N-terminally tagged GFP-αE-catenin vectors were prepared using a lentiviral system^73^ with lentivirus packaging and envelope plasmids (psPAX2 #12260 and pMD2.G #12259, Addgene). Antibodies, DNA and other reagents are listed in the Key Resources Table. All constructs listed will be submitted to Addgene with detailed vector maps per NIH guidelines.

### Cell culture, stable cell line selection and immunoblotting

MDCK II cells were maintained in Dulbecco’s Modified Eagle’s Medium (DMEM, Corning), containing 10% fetal bovine serum (FBS, R&D Systems), 100 U/mL penicillin and 100 μg/mL streptomycin (Corning). MDCK Afadin KO cells were acquired from Tetsuhisa Otani and Mikio Furuse^95^. Our in-house cell line was recently authenticated as canine by RNA sequencing analyses relevant to a different project. α-cat/*Ctnna1* knockout MDCK cells were generated using CRISPR-Cas9 system as described in^73^. α-cat^KO2.2^ MDCK were restored with wild-type and mutant α-cat forms by lentiviral infection, selected in puromycin (5μg/μL)^73^. GFP-α-cat-positive cells were flow sorted (FACSMelody 3-laser sorter (BD)) to ensure even expression across constructs. Cells were lysed in buffer containing 50 mM Tris, pH 7.5, 5 mM EDTA, 150 mM NaCl, 5% glycerol, and 1% Triton X-100 with protease inhibitor cocktail (Roche, Nutley, NJ). For immunoprecipitations, lysates were incubated with indicated antibodies and ImmunoPure Immobilized Protein G or Protein A (Pierce, Rockford, IL). Precipitated proteins were washed and subjected to SDS–PAGE and Western blot analysis using standard procedures. Blots were imaged digitally using a Li-Cor Odyssey blot imager and quantified using Li-Cor Image Studio software (Li-Cor Biosciences). Primary and secondary antibodies used are provided in the Key Resources Table.

### Immunofluorescence and Imaging

Cells were grown on cell culture inserts (BD Falcon 353494; high density 0.4μm) for 10+ days), fixed in 4% paraformaldehyde (Electron Microscopy Services, Hatfield, PA) for 15’, quenched with glycine, permeabilized with 0.3% Triton X-100 (Sigma) and blocked with normal goat serum (Sigma). Cells were alternatively cultured ∼48 h and imaged on glass coverslips (Figure S5. Primary antibody incubations were performed at RT for 60’ or 4°C overnight; secondary incubations at RT for 30’, interspaced by multiple washes in PBS. Coverslips were mounted in ProLong Gold fixative (Life Technologies). Fixed images of GFP-α-cat, Afadin and F-actin localizations were captured with Nikon A1R Confocal Laser Point Scanning microscope using NIS Elements software (Nikon) with GaAsP detectors and equipped with 95B prime Photometrics camera (Photometrics), Plan-Apochromat 60x/1.4 objective (Figures 1, 4, 6). Fixed images of GFP-α-cat, Afadin, and F-actin localizations were captured with Nikon AXR Confocal Laser microscope with widefield resonant GaAsP detectors and equipped with a sCMOS camera (Photometrics), Plan-Apochromat 60x/1.4 objective (Figures S6, S7). Confocal Z-stacks were taken at step size of 0.25μm. RGB histology images were acquired using a Nikon Ti2 Widefield microscope with a Nikon DS-Qi2 camera, LiDa light source (Lumencor), Plan-Apochromat 4x/0.2 objective (Fig. 7), or using a TissueGnostics automated slide imaging system (replicates, not shown).

### Image analysis and fluorescence quantification

Junctional enrichment of afadin, α-cat, or F-actin was quantified as previously described (Quinn et al., 2024). Briefly, raw integrated signal intensity of 1um circular ROIs was measured on bicellular junctions (BCJ) and adjacent cytoplasm (Cyto) for each channel. Cyto signal was subtracted from BCJ signal, and the enrichment was normalized. Measurements were performed on 1.5um maximum intensity projections centered at the maximum apical α-cat signal. To assess the apical distribution of α-cat signal in Fig. 1E, z-stacks were analyzed for the slice with the apical most α-cat signal. This z-slice position was set to 0 to center apical junction depth across replicate stacks. The α-cat signal intensity for each z-slice was normalized to the total % α-cat signal, and the average signal for each experimental group was plotted by depth. In Fig. 4C, intensity projections were made from z-stacks to include all/basal signal, signal from the 1.5um apical junction, or a single slice at the maximum apical α-cat signal. Colocalization was performed on these images in a FIJI plugin (JaCoP)^96^, using the default Manders’ 1 settings to report α-cat:Afadin overlap fraction. In Fig. 6C, monolayer visual defects were manually quantified by counting the total number of junctions and junction-level breaks in z-stacks, calculating the frequency of breaks by vertex number, and averaging frequencies across replicate FOVs. In Fig. S5D, the relationship of α-cat and Afadin was plotted using circular ROIs as above. The junction intensity from the two channels was normalized, plotted, and was analyzed using GraphPad’s Pearson Correlation function. To quantitatively compare hydrocephalus severity (Fig. 7E), imaged mouse brain sections were traced using FIJI’s polygon selection tool. Ventricular spaces were outlined, and the measured free area was divided by the total brain area. Replicates were assigned ‘Anterior’ or ‘Posterior’ depth based on Nissl-staining patterns and the Allen Mouse Brain Atlas. Data statistical analysis was done using GraphPad and detailed in figure legends for each experiment.

### In vitro binding site mapping and kinase assay

Pulldown was performed using indicated αE- or αN-catenin forms, and interaction strength was assessed by Coomassie stating. The GST-fusion pulldown construct containing the rat Afadin residues 1391-1449; MDLPLPPPPANQAAPQSAQVAAAERKKREEHQRWYEKEKARLEEERERKRREQERKL GQ was subcloned into pGEX4T1 using BamHI/XhoI sites. The original rat Afadin cDNA was a gift from the Takai lab. Residues in the putative Afadin-binding region of α-cat were subject to NMR titration, with several residues exhibiting a broadening (strong) chemical shift. These residues clustered together at the surface highlighted in Fig. 5C. Specific residues were mutagenized and subjected to Afadin pulldown. The identified residues were used for *in silico* and *in situ* experiments. In vitro kinase assays were carried out as previously described^66^.

### Protein Modeling

AlphaFold2 in ColabFold^97^ and MDOCKPP^98^ were used to generate 3D models of α-cat. AlphaFold predictions were ran using unsupervised or template-guided inputs (α-cat PDB structures: 4IGG, 1H6G, 7UTJ, 6WVT, 1L7C, 4K1N). The predicted interactions of WT or Δ-Afdn α-cat with afadin was modeled using the sequence of the identified α-cat M3 helix1 and Afadin residues 1410-1450. Protein structures were visualized from pdb files in PyMol. Multiple sequence alignments of α-cat by isoform and species were performed using FastA files downloaded from NCBI’s BLAST and aligned with Clustal Omega in EMBL-EBI’s sequence analysis Job Dispatcher^99^.

### TEER and Paracellular Flux Assays

Cells were grown on culture inserts as above for 10 days, after which barrier was assessed with transepithelial electrical resistance (TEER) or a fluorescence flux assay. For TEER (Fig. 6E), monolayer and background resistance was measured using a EVOM2 Voltometer with a STX2 probe (World Precision Instruments). TEER was calculated by subtracting the blank value and multiplying by the filter area. Normalized values and averages were plotted in Prism. To measure flux (Fig. 6F), cells were washed and held in supplemented phenol-red free DMEM. The top chamber was supplemented to contain 1mg 4-kD FITC-dextran (Sigma). Cells were returned to the incubator, and aliquots of media from the basal compartment were taken at timed intervals. Fluorescence leak was determined using a SpectraMax M2e plate reader equipped with SoftMax Pro 6.4 software (Molecular Devices). Background intensity of a measured blank was subtracted, and normalized fluorescence was plotted in Prism.

### Mouse model generation by CRISPR-mediated gene editing

We introduced 4 codon mutations (S641A, S652A, S655A, T658A) into *Ctnna1* exon 14 using CRISPR gene editing, creating the *Ctnna1-P-linker (Ctnna1*^4A^) mouse. Two separate experiments were required to generate *Ctnna1*^4A^ F0 founders. In the first experiment, CRISPR components were microinjected into the cytosol of fertilized C57Bl/6J embryos along with an exogenous repair template encoding the 4 codon mutations (sequences below). Editing was successful; however, no founders with all 4 mutations were generated. One F_0_ male harbored 3 of the 4 mutations (CK1 sites, S652A, S655A, T658). This mouse was breed with a C57BL/6J females to establish the line and offspring were then bred to homozygosity for the 3 *Ctnna1* mutations. Homozygous males and het- or homozygous females were used for subsequent crosses. To create the 4^th^ mutation (CK2 site, S641A), we electroporated the CRISPR components and a new repair template (specific for the 4^th^ mutation) into fertilized embryos from edited mice. Two founders were born and genotyped for editing events via PCR, gel electrophoresis and Sanger sequencing. Mosaic F_0_ mice harbored an allele with the 4^th^ mutation. These mice were validated for germline transmission of the entire 4 codon mutation sequence (S641A, S652A, S655A, T658A), generating two *Ctnna1*^4A/WT^ founder lines, 1034 and 1035. Founder lines were backcrossed to C57BL/6 mice for 10 generations, developing two congenic strains to dilute phenotypic contributions from possible off-target effects of CRISPR guides. Figure 7 documents genotypic frequencies at the time of weaning and incidence of developmental defects across the allelic series (*Ctnna1*^WT/WT^; *Ctnna1*^4A/WT^; *Ctnna1*^4A/4A^). Genotyping is routinely determined by a validated qPCR-based system (Transnetyx). Mouse tissue protein analysis, organs were homogenized in RIPA buffer with protease and phosphatase inhibitor mini tablets (Pierce) using a Tissue-Tearor (985370-395, BioSpec Products). All experimental protocols were approved by the Institutional Animal Care and Use Committee at Northwestern University (Chicago, IL, USA). All strains were bred and housed at a barrier- and pathogen-free facility at the Center for Comparative Medicine.

### gRNA/Cas9 design, expression and mouse line availability

We used 2 gRNA to target the 5’ and 3’ regions of *Ctnna1* exon14; gRNA2 targets the 5’ region (5’– GCTGCAGACCCCCGAGGAGT-3’); gRNA1 targets the 3’ region (5’-TCAGCTGGTCATCTTCTGTC-3’) (Fig. 7A). gRNA was cloned into the pX458 Cas9 vector (Addgene #48138) for in vitro transcription (MEGAshortscript T7 transcription kit (Thermo, AM1354). gRNA transcripts were combined with Cas9 mRNA (Thermo, A29378) and delivered to the cytosol of fertilized C57Bl/6J embryos along with an exogenous repair template encoding codon mutations. Detailed write up from Northwestern’s Transgenic and Targeted Mutagenesis Laboratory with full sequences is available upon request. The mouse line is currently cryopreserved at Northwestern and The Jackson Laboratories.

## KEY RESOURCES TABLE

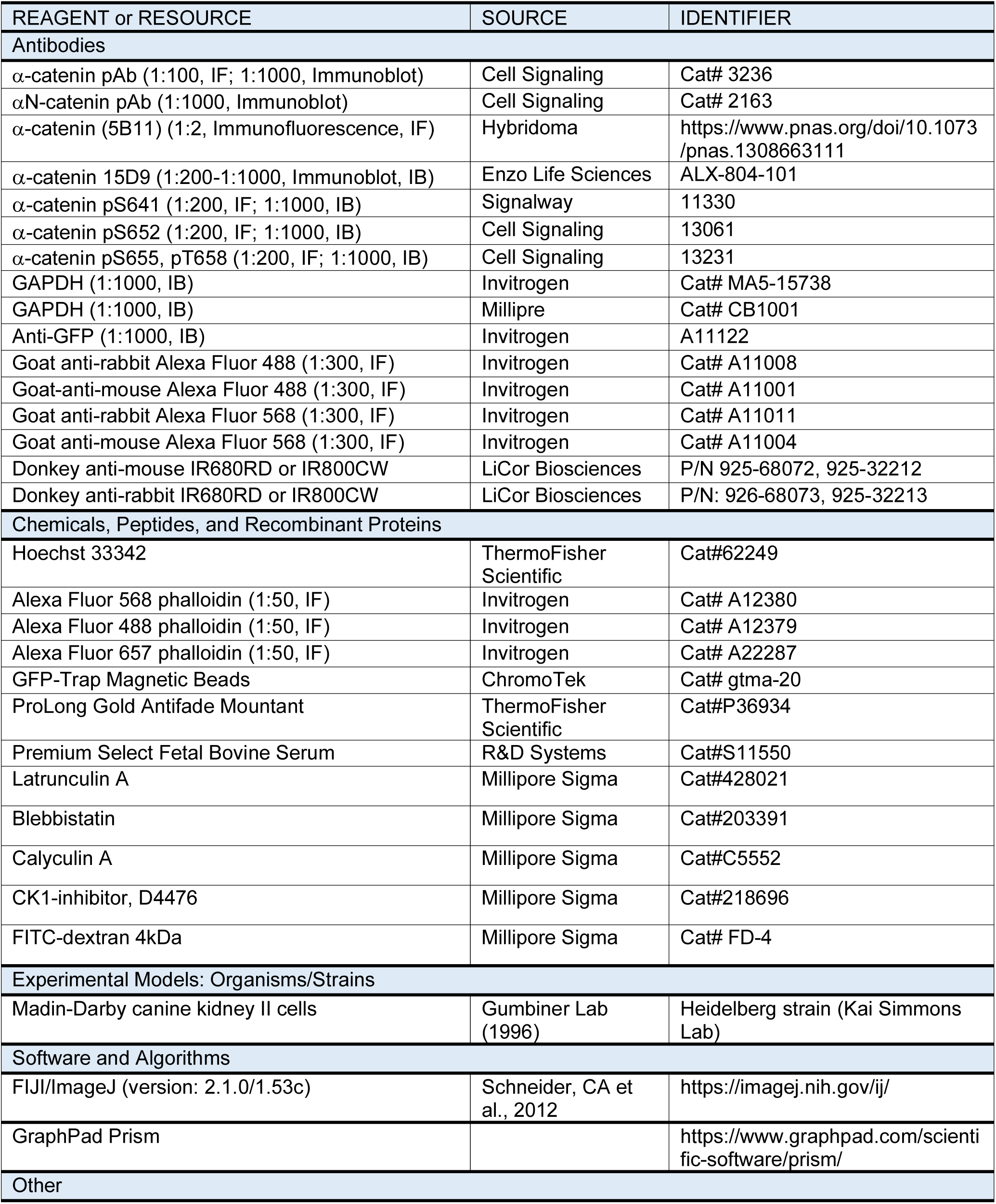
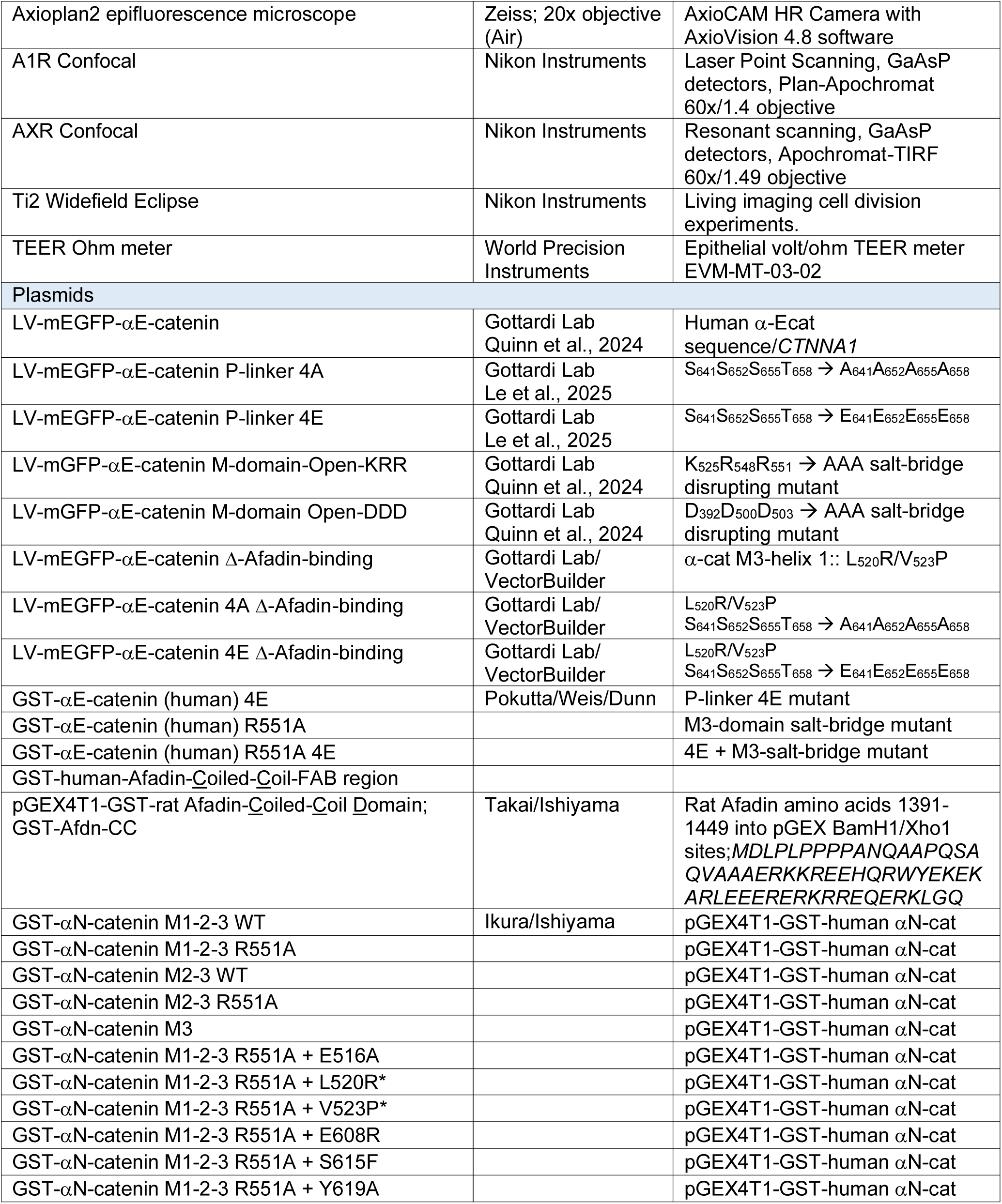

## ACKNOWLEDGEMENTS

We thank Tetsuhisa Otani (Tokyo Metropolitan University) for sharing Afadin-KO MDCK cells. This work relied on the following Northwestern University services and core facilities: Center for Advanced Microscopy (NCI CCSG P30 CA060553, NCRR 1S10 RR031680, 1S10OD021704), Flow Cytometry (NCI CA060553, 1S10OD011996, 1S10OD026814) and Transgenic and Targeted Mutagenesis Laboratory (NCI CA060553).

## Author Contributions

**Jeanne M. Quinn:** Investigation, Methodology, Formal Analysis, Writing (original draft)**, Phuong M. Le:** Investigation, **Anthea Weng:** Investigation, **Annette S. Flozak:** Investigation, Formal Analysis, Writing (review and editing), **S. Sai Folmsbee:** Investigation, **Erik Arroyo-Colon:** Formal analysis, **Mitsu Ikura:** Conceptualization, Acquisition of Funding, **Noboru Ishiyama:** Investigation, Conceptualization, **Cara J. Gottardi:** Conceptualization, Supervision, Acquisition of Funding, Writing (original draft, review and editing).

---

**Figure S1:**
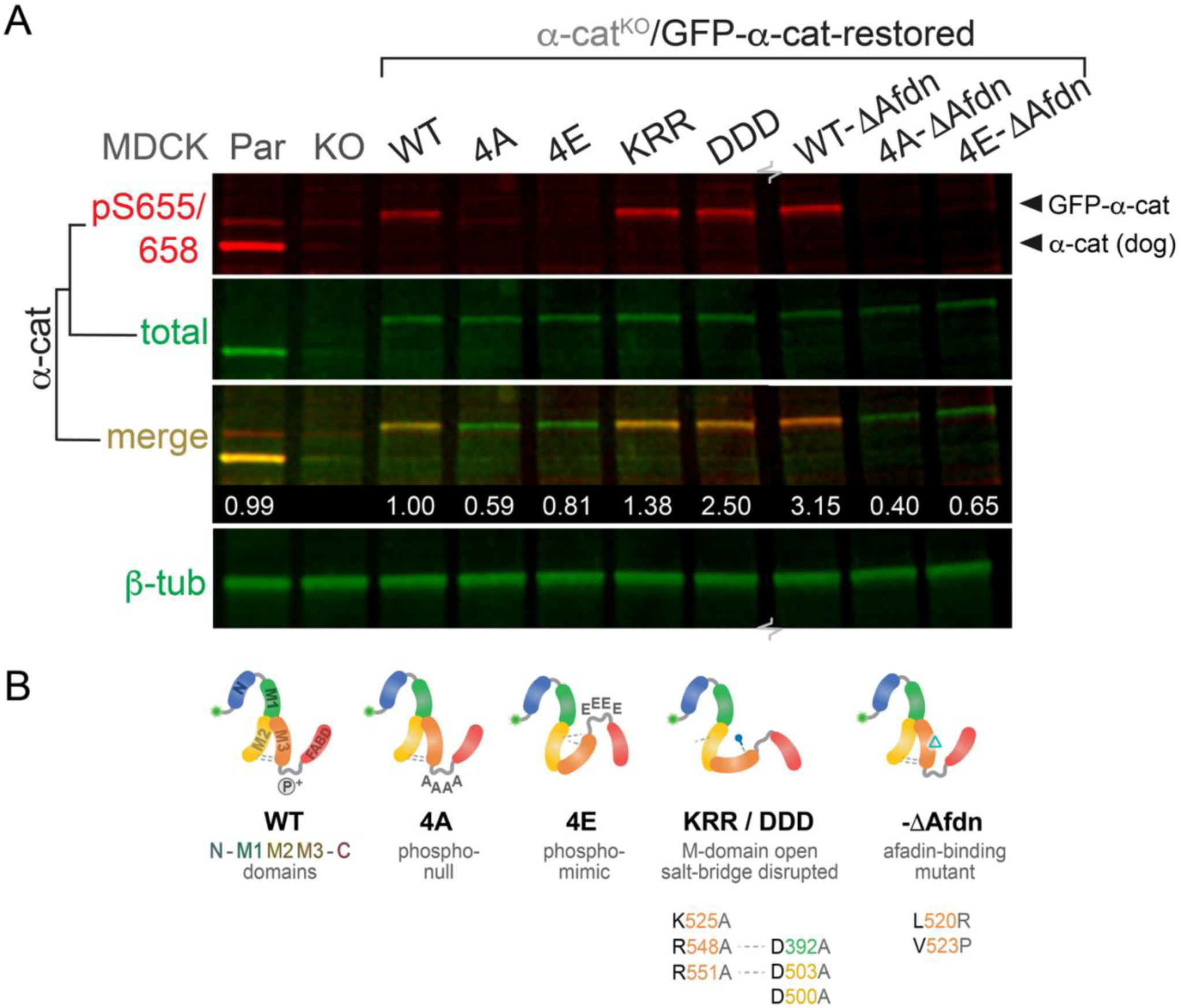
Restoration of GFP-α-cat mutants in α-cat CRISPR-KO MDCK cells. (**A**) Immunoblot of MDCK cells restored with GFP-tagged forms of α-cat via lentiviral delivery. Total-α-cat antibody detects similar α-cat expression across restored cell lines. Specific phospho-sites removed in 4A (phospho-mutant) and 4E (phospho-mimic) are not detected by the phospho-specific antibody. Ratio of phospho-to-total α-cat was quantified and standardized to WT. β-tubulin was used as loading control. α-cat restored lines were sort-matched by flow cytometry and periodically blotted to ensure against expression drift. (**B**) Schematic of GFP-α-cat mutant forms used in this study.

**Figure S2:**
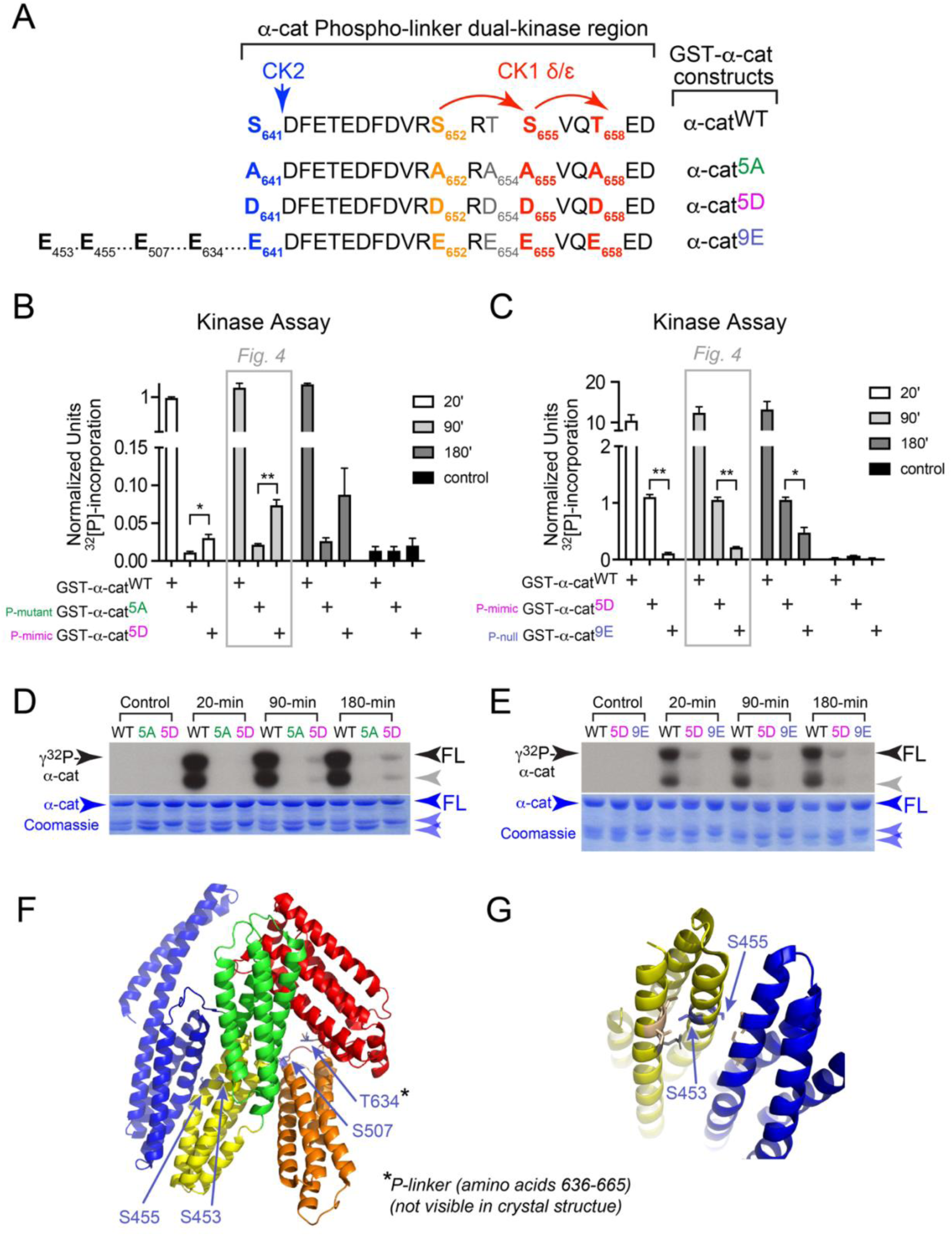
α-cat P-linker phospho-mimic promotes M-domain accessibility via in solution kinase assay. (**A**) Schematic of α-cat constructs with phospho-proteomic defined P-linker S/T residues replaced by non-modifiable phospho-mimic (aspartate, (D) or glutamate (E)) and phospho-mutant (alanine, (A)) residues. We previously showed that CK2 phosphorylates α-cat at S641, whereas CK1 sequentially phosphorylates α-cat at S652, S655 and T658 (Escobar et al., 2015). For loss-of-phosphorylation studies, T654 was also mutated, because pT654 could not be confidently distinguished from pS655 by mass spectrometry (Harvard Taplin Facility). Since CK1 typically phosphorylates proteins at S/TxxS/TxxS/T residue spacing, we favor the phospho-scheme shown in A (red arrows), but it is formally possible T654 is modified non-canonically. Thus, these early in vitro phosphorylation studies relied on P-linker 5-residue mutant (α-cat-5A, green) and mimic proteins (α-cat-5D, magenta), respectively. An α-cat 9-residue total phospho-mutant (α-cat 9E or P-null, blue), prevented modification of 4 additional phospho-sites localizing to α-cat’s M-domain (S453, S455, S507 and T634). These latter sites were identified in our original phospho-proteomic mass spectrometry analysis of α-cat^66^ (Harvard Taplin Facility). (**B-E**) Kinase activity is measured by the incorporation of γ^32^P-incorporation over time. After incubation, GST-purified α-cat was subjected to gel electrophoresis, Coomassie staining, gel drying and exposure to film. (**D**-**E**) Kinase accessibility was quantified (**B-C**) by standardizing the full-length (FL) γ^32^P-containing α-cat bands (film exposure, grayscale image) to total α-cat detected by Coomassie stain (lower image, purple). Normalized values from 3 experiments were plotted and compared by multiple t-tests; *P<0.05, **P<0.01. (**F-G**) α-cat M-domain residues mutagenized in construct P-null 9E (blue). (**F**) Full length α-cat crystal structure highlighting locations of S/T residues in M2 and M3. All mutagenized S/T residues were evaluated for potential confounding salt-bridge interactions with nearby residues. (**G**) Zoomed-in view of the residues in M2, with nearby sidechains showing no charge-based interactions. Evidence these residues (S453, S455, S507 and T634) are more accessible to in vitro phosphorylation in the α-cat 5E versus α-cat 5A mutant suggests that α-cat P-linker modification allosterically alters the M-domain.

**Figure S3:**
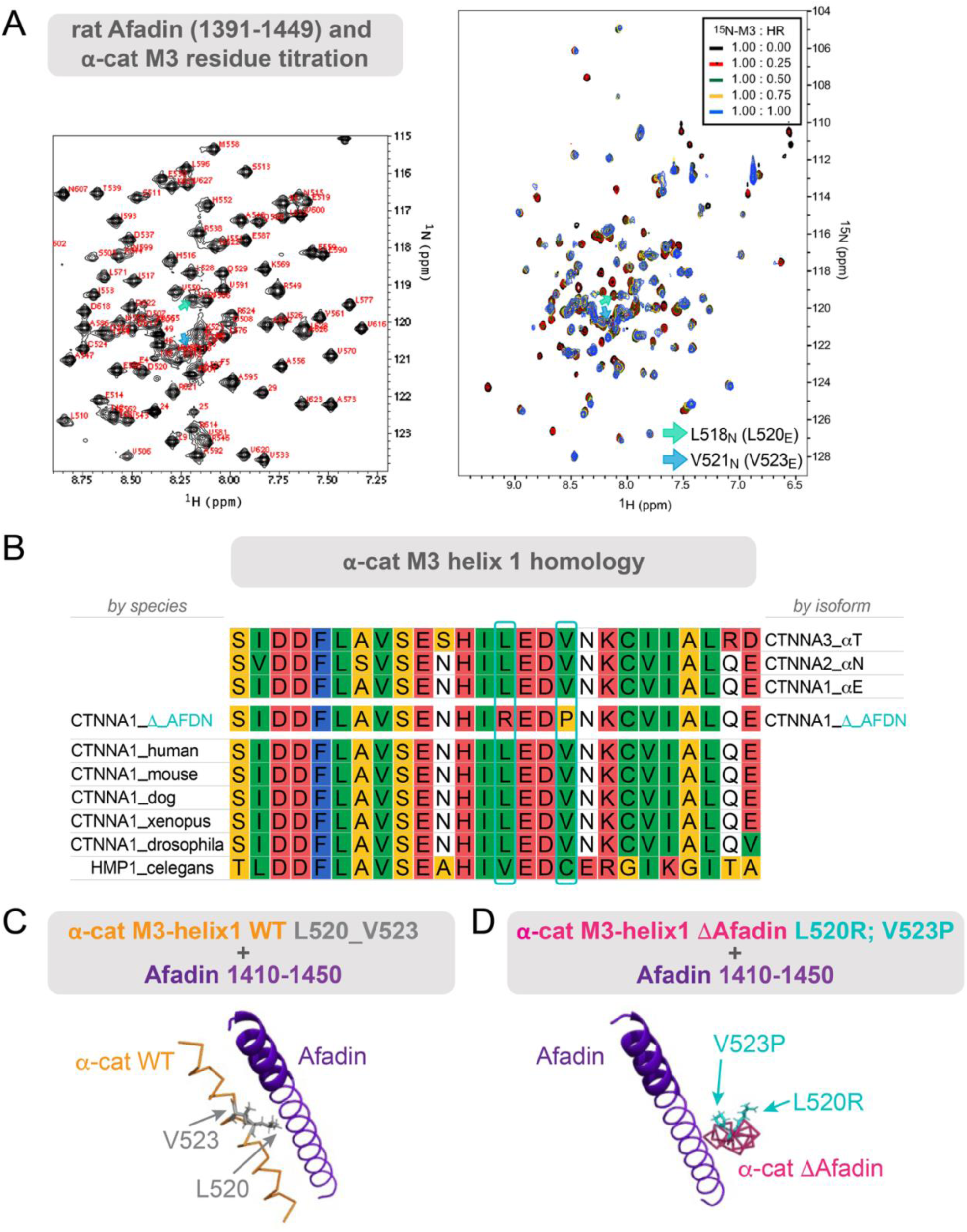
Modeling of the α-cat M3/Afadin-binding interaction. **A**) NMR titration analysis of ^15^N-labeled αN-cat-M3 interaction with unlabeled Afadin-CC-ABR. Overlay of ^1^H-^15^N HSQC spectra of ^15^N-αN-cat-M3 with unlabeled Afadin-CC-ABR at various molar ratios are shown, indicating Afadin binding by select αN-cat-M3 residues. These data were used to validate the binding interface in Fig. 5C. **B**) Multiple sequence alignment of helix 1 in αE-cat (*CTNNA1*) M3-domain demonstrates preserved residues across species and isoforms. **(C-D)** Predictions from molecular docking simulations between Afadin and α-cat WT or ΔAfdn. (**C**) α-cat M3 helix 1 is predicted to interact with Afadin through residues L520 and V523. (**D**) Afadin and α-cat ΔAfdn docking predictions yield misaligned interactions.

**Fig. S4:**
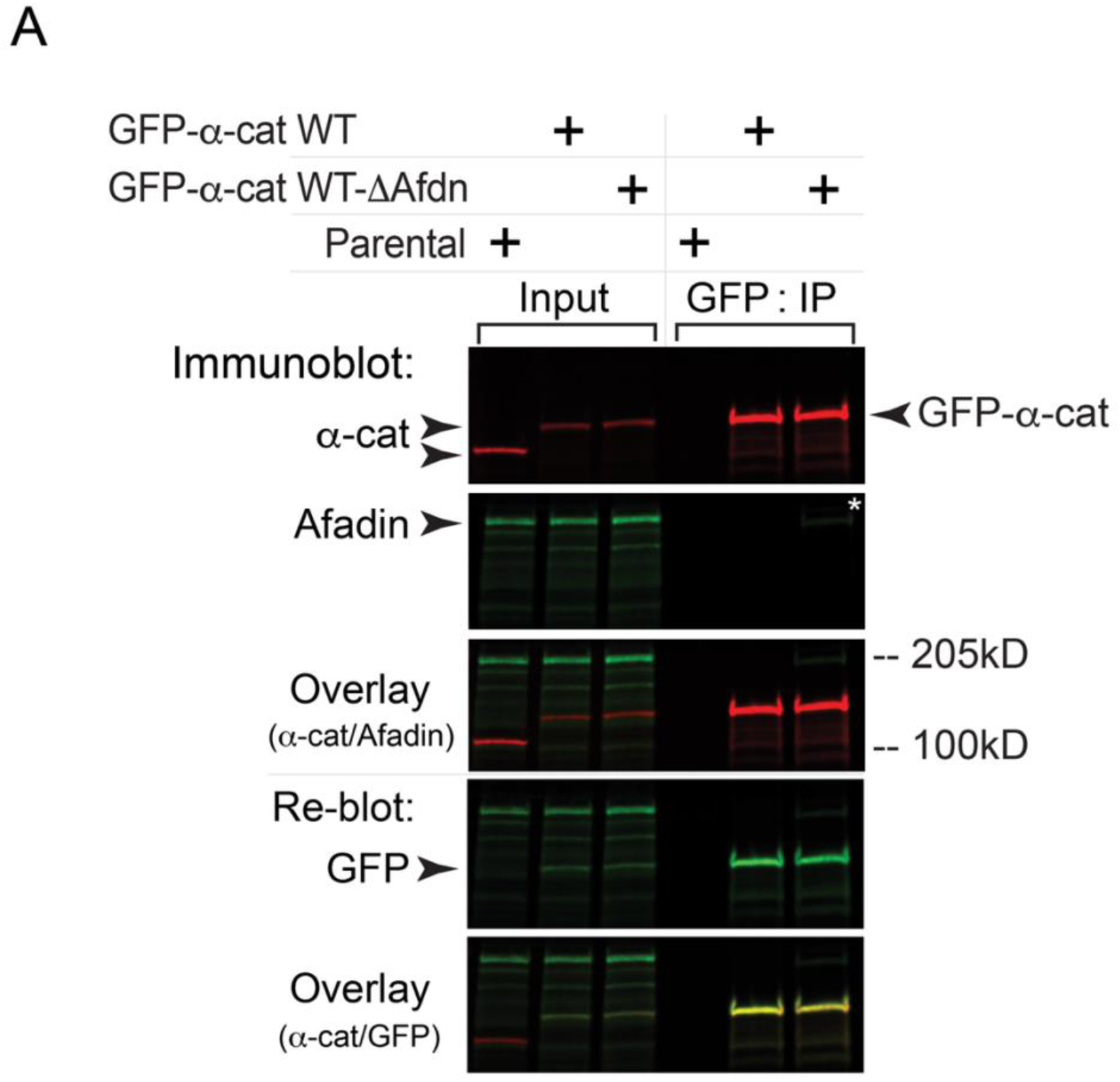
Afadin does not substantially co-immunoprecipitate with α-cat in MDCK epithelial cell lysates. Immunoblot validation of α-cat WT and ΔAfadin (Afdn) protein expression in α-cat CRISPR KO MDCK cells. Parental MDCK cells shown (left-most lane). Note that despite robust GFP-α-cat affinity precipitation in both cell lines, Afadin co-immunoprecipitation was weak.

**Figure S5:**
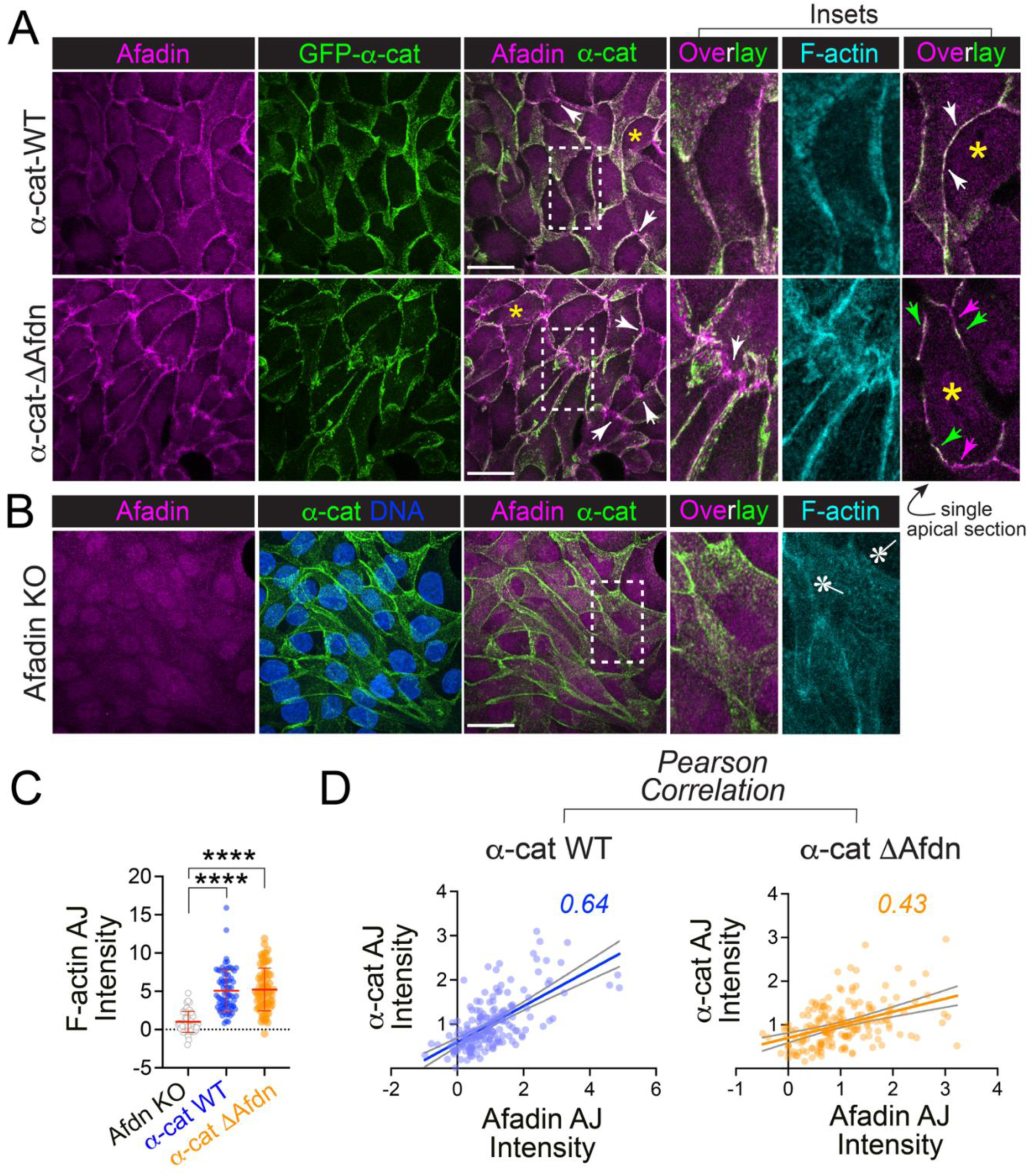
α-cat M3-Δ-Afadin binding mutant does not phenocopy Afadin KO MDCK cells matured on glass. (**A**) Confocal images of glass coverslip plated MDCK monolayers with restored GFP-α-cat WT and Δ-Afdn forms fixed and immuno-stained for Afadin (magenta) and F-actin (phalloidin, cyan). Native GFP-α-cat is shown in green. *En face* images are maximum intensity projections (M.I.P.). Scale bar = 20μm. Dotted boxes correspond to expanded inset views (right). Arrows point to multi-vertex junctions that appear to show less α-cat/Afadin colocalization for the α-cat Δ-Afdn mutant. This is especially clear for the apical most portion of bi-cellular junctions (single apical section), where white arrows show better colocalization between α-cat WT and Afadin versus the α-cat Δ-Afdn mutant, where separation of α-cat (green arrows) and Afadin signal (magenta arrows) is shown. Asterisk (yellow) shows corresponding cell in lower mag view; note that views are rotated. (**B**) Confocal images (maximum intensity projection) of Afadin KO MDCK cells grown on glass. Dotted box corresponds to expanded insets (right). Note that F-actin recruitment to cell-cell junctions is reduced in Afadin KO cells (asterisks), as previously shown^15^. Note F-actin recruitment is not obviously altered between α-cat WT and α-cat Δ-Afdn mutant cells (A, above insets). (**C**) Quantification of F-actin junction intensity from images in A and B, using 1μm circular ROIs taken from bicellular junctions, subtracting adjacent cytoplasm signal. Normalized fluorescence was plotted, with symbols corresponding to 1 biological experiment analyzing 50-75 junctions/construct. ****P<0.0001 by ANOVA with Tukey’s multiple comparison. (**D**) Quantification of α-cat/Afadin colocalization differences seen in A across multiple junctions. Intensity correlation was analyzed by Pearson’s (WT r = 0.6430, ****p < 0.0001; α-cat Δ-Afdn r = 0.4298, ****p < 0.0001). This suggests weaker colocalization between α-cat Δ-Afdn mutant and Afadin at zAJs than α-cat WT and Afadin.

**Figure S6:**
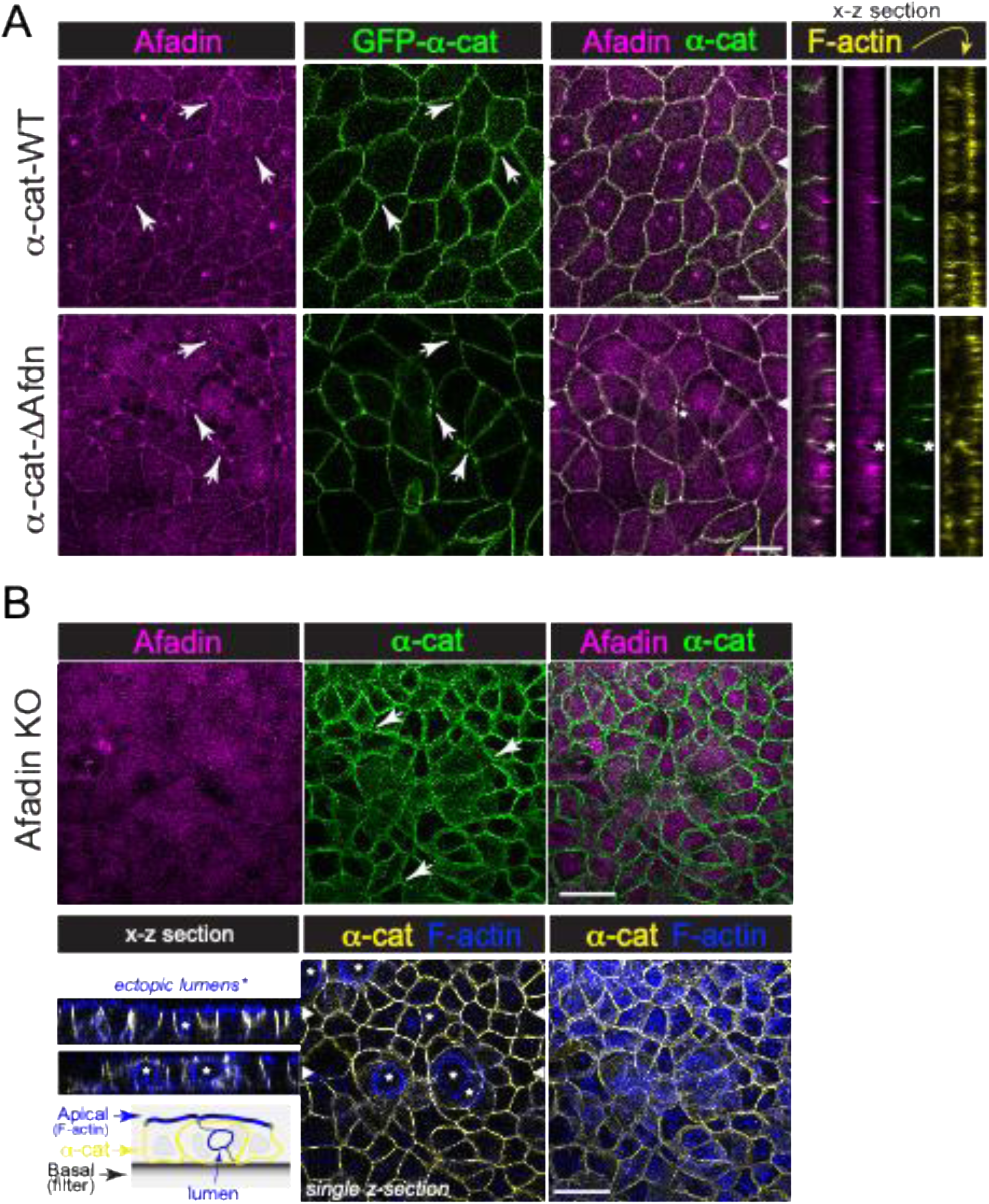
α-cat M3-Δ-Afadin binding mutant does not phenocopy Afadin KO MDCK cells matured on filters. (A) Confocal images of MDCK monolayers matured on filters with restored GFP-α-cat WT and Δ-Afdn forms fixed and immuno-stained for Afadin (magenta) and F-actin (phalloidin, yellow). Native GFP-α-cat is shown in green. *En face* images are single x-y optical slices. White arrows show multi-vertex junctions, which clear gaps for the α-cat Δ-Afdn mutant. Scale bar = 5μm. Orthogonal (x-z) views are shown to right; section marked by white arrowheads in overlay image. Asterisk shows recessed apical junctions extending basally in the α-cat Δ-Afdn mutant. (**B**) Confocal images (maximum intensity projection) of Afadin KO MDCK cells matured on filters (2-weeks), fixed and immuno-stained for Afadin (magenta). Arrows show normal (un-recessed) multi-vertex junctions. Scale bar = 20μm. Note that on filters, we did not detect an obvious loss of F-actin recruitment to zAJ (F-actin, blue; α-cat, yellow), although Afadin KO cells showed greater capacity to form intra-epithelial (ectopic) lumens (lower left, schematic).

**Figure S7:**
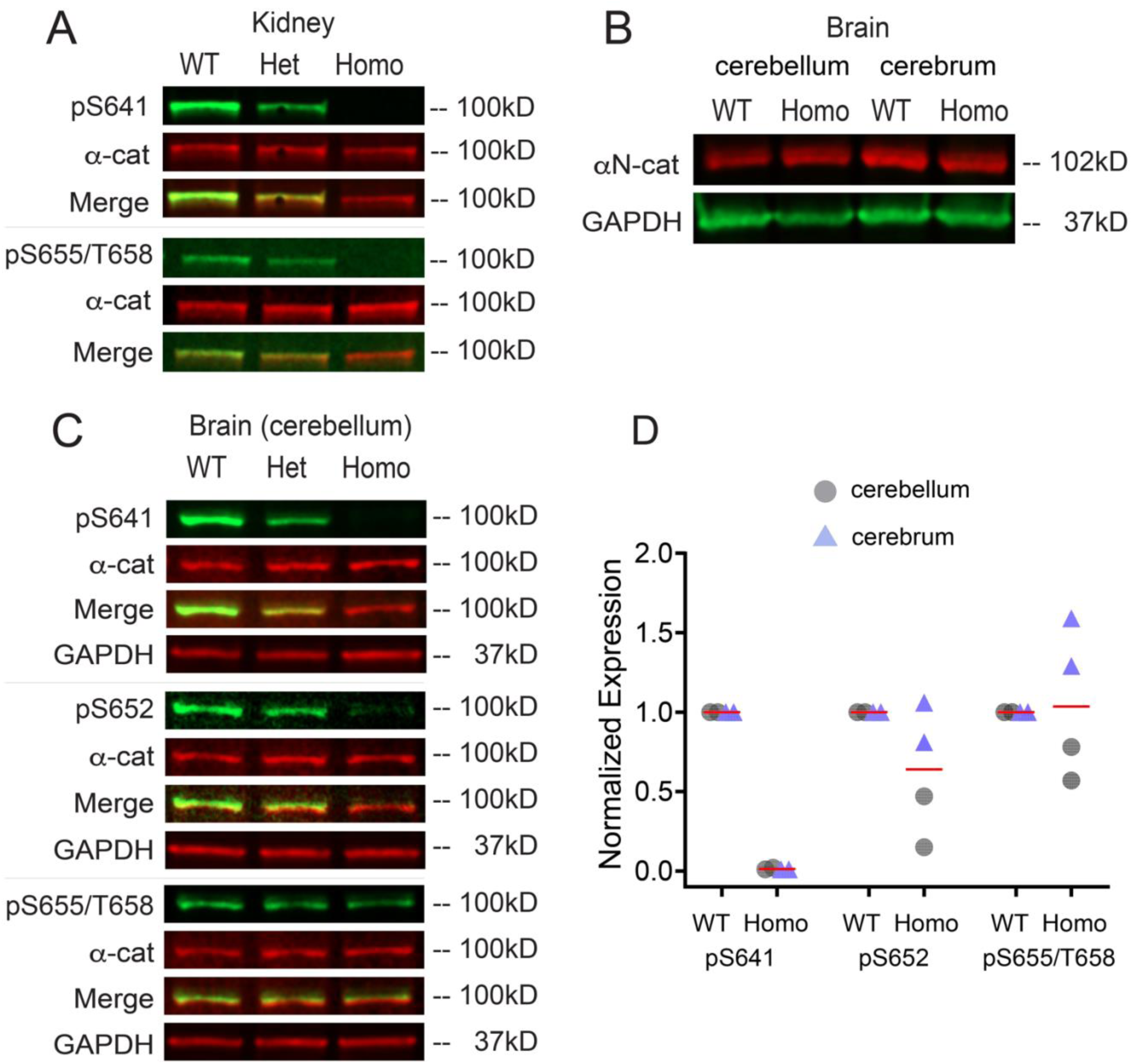
Immunoblot validation of *Ctnna1*-phospho-mutant knock-in mouse line by tissue. (**A**) Lysates prepared from *Ctnna1*^WT/WT^ (Wild-type, WT), *Ctnna1*^WT/4A^ (heterozygous, HET) or *Ctnna1*^4A/4A^ (homozygous, HOMO) mouse kidney show that α-cat phosphorylated at S641 or S655/T658 are reduced or ablated in kidney lysates prepared from HET or HOMO mutant mice relative to a littermate WT control. Immunoblot detection using a LiCOR imaging system where nitrocellulose was simultaneously incubated with antibodies to phospho-α-cat (green) or total α-cat (red) (**B**) Lysates prepared cerebellum corresponding to cerebrums analyzed in Figure 7 and subjected to immunoblotting as in A. While detection of α-cat phosphorylated at S641 is completely blocked in this tissue, pS652 is reduced and pS655/pT658 immunodetection persists in the *Ctnna1*^4A/4A^ (HOMO) condition. Since brain (cerebellum and cerebrum) expresses the neural isoform of α-cat, *Ctnna2*/αN-catenin, we affirmed expression of this isoform in C. (**C**) Immunoblot with antibody to αN-cat, showing abundant expression of this protein. (**D**) Graph quantification of α-cat pS641, pS652 and pS655/pT658 detection from brain lysates (cerebellum and cerebrum) of two different animals. Collectively, these data show that the *Ctnna1*^4A/4A^ HOMO-knock-in mouse completely blocks phosphorylation of αE-cat pS641, pS652 and pS655/pT658 in tissues that only express αE-cat/Ctnna1 (A, kidney). Tissues that express other α-cat isoforms, such as brain, reveal detection of identical phospho-sites in these isoforms (αN-cat; αT-cat).

